# The population genomics of transposable element activation in the highly repressive genome of an agricultural pathogen

**DOI:** 10.1101/2020.11.12.379651

**Authors:** Danilo Pereira, Ursula Oggenfuss, Bruce A. McDonald, Daniel Croll

**Affiliations:** Plant Pathology, Institute of Integrative Biology, ETH Zürich, Zürich, Switzerland; Laboratory of Evolutionary Genetics, Institute of Biology, University of Neuchâtel, Neuchâtel, Switzerland

**Keywords:** adaptation, genetic diversity, selfish elements, pathogen, transposable elements, repeat-induced point mutations

## Abstract

The activity of transposable elements (TEs) can be an important driver of genetic diversity with TE-mediated mutations having a wide range of fitness consequences. To avoid deleterious effects of TE activity, some fungi evolved highly sophisticated genomic defences to reduce TE proliferation across the genome. Repeat-induced point (RIP) mutations is a fungal-specific TE defence mechanism efficiently targeting duplicated sequences. The rapid accumulation of RIP mutations is expected to deactivate TEs over the course of a few generations. The evolutionary dynamics of TEs at the population level in a species with highly repressive genome defences is poorly understood. Here, we analyze 366 whole-genome sequences of *Parastagonospora nodorum*, a fungal pathogen of wheat with efficient RIP. A global population genomics analysis revealed high levels of genetic diversity and signs of frequent sexual recombination. Contrary to expectations for a species with RIP, we identified recent TE activity in multiple populations. The TE composition and copy numbers showed little divergence among global populations regardless of the demographic history. Miniature inverted-repeat transposable elements (MITEs) and terminal repeat retrotransposons in miniature (TRIMs) were largely underlying recent intra-species TE expansions. We inferred RIP footprints in individual TE families and found that recently active, high-copy TEs have possibly evaded genomic defences. We find no evidence that recent positive selection acted on TE-mediated mutations rather that purifying selection maintained new TE insertions at low insertion frequencies in populations. Our findings highlight the complex evolutionary equilibria established by the joint action of TE activity, selection and genomic repression.

**Data Summary:** All Illumina sequence data is available from the NCBI SRA BioProject numbers PRJNA606320, PRJNA398070 and PRJNA476481 (https://www.ncbi.nlm.nih.gov/bioproject). The Methods and Supplementary Figures S1-S11 and Supplementary Tables S1-S4 provide all information on strain locations and outcomes of genome analyses.

## Introduction

Genetic diversity in natural populations largely determines the evolutionary potential of populations. Polymorphism and diversity in haplotypes can arise from various processes including single base mutations (Baranova et al. 2015), reshuffling of alleles through recombination (Goddard et al. 2005), chromosomal rearrangements (Wolfe & Shields 1997) and the action of selfish DNA sequences (Horns et al. 2017). Transposable elements (TEs) are ubiquitous selfish elements capable of proliferating throughout the genomic landscape (Kidwell 2002). Transposition activity can increase with senescence (De Cecco et al. 2013), or under environmental stress conditions increasing the risk for genetic modifications (Chen et al. 2015). TE-mediated genetic changes can impact fitness of the host. During transposition, TEs can create genetic variation by altering coding sequences, gene regulation, or triggering chromosomal rearrangements through non-homologous recombination (Hedges & Deininger 2007; Burns & Boeke 2012). The impact of TEs is in most cases negative or neutral, but can rarely also be positive by contributing to adaptation in humans and other organisms (Chou et al. 2002; Desalvo et al. 2008). In plant pathogens, TEs are important drivers of genome evolution and adaptation to the host (Raffaele & Kamoun 2012; Möller & Stukenbrock 2017; Seidl & Thomma 2017). For example, TE activity caused gene deletions and sequence reshuffling in the fungal pathogen *Blumeria graminis* f. sp *hordei* ultimately underpinning host specialization (Spanu et al. 2010). In *Zymoseptoria tritici*, gain in virulence was observed after TE-mediated deletion of a single gene (Hartmann et al. 2017), and fungicide resistance emerged from overexpression and splicing alterations of target genes after upstream TE insertions (Omrane et al. 2015; Steinhauer et al. 2019). Despite the major impact on genome stability and the expression of phenotypic traits, ongoing TE activity at the intra-specific level is poorly understood in plant pathogens and other organisms. Major questions remain regarding the genome-wide distribution of active TEs and their impact on fitness.

A number of fungal genomes, including the genomes of various plant pathogens, contain regions enriched in TEs and are evolving at faster evolutionary rates largely through sequence rearrangements (Raffaele & Kamoun 2012; Dutheil et al. 2016; Faino et al. 2016; Wang et al. 2017). Polymorphism in genes located within such regions is usually higher than in more conserved regions (Rouxel et al. 2011; Wang et al. 2017). Furthermore, key virulence factors tend to localize in the vicinity of repetitive regions (Möller & Stukenbrock 2017; Fouché et al. 2018; Richards et al. 2019). Genes showing evidence for presence-absence polymorphism were shown to be closer to TEs than conserved genes (Yoshida et al. 2016; Richards et al. 2019; Hartmann & Croll 2017). Gene deletions in some pathogens are associated with gains in virulence if the affected proteins are recognized by the host (Hartmann & Croll 2017). Controlling the activity of TEs faces costly trade-off between genomic defence mechanisms such as silencing and the *bona fida* expression of nearby genes (Hollister & Gaut 2009). Suppression of TE proliferation can be mediated by heterochromatin modifications, DNA methylation, meiotic silencing, and posttranscriptional regulation (Nolan 2005; Gladyshev 2017). A powerful fungal-specific mechanism is to hypermutate duplicated sequences through repeat-induced point (RIP) mutations, a mechanism first described in *Neurospora crassa* (Selker 1990). During sexual recombination, RIP acts via homology recognition changing C:G nucleotides to T:A nucleotides, leaving sequence hallmarks found in genomes of many plant pathogenic fungi (Ikeda et al. 2002; Hane & Oliver 2008; Fudal et al. 2009; Dhillon et al. 2014; Van de Wouw et al. 2019). Although genomic defenses against TEs are frequent across fungi, substantial variation in TE abundance was found among genomes between species but also within species (Plissonneau et al. 2018; Syme et al. 2018; Wyatt et al. 2018; Badet et al. 2020). The population frequency of individual TE sequences is restrained by purifying selection (Oggenfuss et al. 2020). A loss of control or neutral effects can permit copy number expansions (Fouché et al. 2020; Oggenfuss et al. 2020). In rare cases, a TE insertion locus can undergo a selective sweep indicating that the insertion may have created adaptive genetic variation. Considering that genomic defences such as RIP were shown to vary between species (Smith et al. 2012; Van de Wouw et al. 2019), the interplay of genomic defenses, selection and demography has the potential to drive or restrain TE proliferation among populations.

In the wheat pathogen *Parastagonospora nodorum*, TEs caused significant genetic alterations and likely facilitated the transfer of a key virulence gene. *P. nodorum* is a pathogen with high evolutionary potential to adapt to local conditions (Richards et al. 2019; Pereira, McDonald, et al. 2020; Pereira, Croll, et al. 2020) and a worldwide distribution causing significant wheat yield losses (Oliver et al. 2012). The pathogen harbors a repertoire of various effector genes (named *Tox* genes) that confer host adaptation by causing cell death on susceptible plants. Interestingly, the effector gene *ToxA* was involved in a horizontal gene transfer (HGT) in the triad of the pathogens *P. nodorum, Pyrenophora tritici-repentis*, and *Biopolaris sorokiniana* (Friesen et al. 2006; McDonald et al. 2019). The HGT was facilitated by the TE environment in which *ToxA* is embedded. Overall, repetitive elements in the genome of *P. nodorum* comprise less than 6.2% of the total genome size (Hane et al. 2007; Richards et al. 2019), proximity to TEs was correlated with gene presence-absence polymorphisms within populations (Richards et al. 2019). Despite extensive evidence for RIP-mediated TE control in the reference genome (Hane & Oliver 2008, 2010), the species display remarkable examples of TE-mediated phenotypic trait variation. Hence, TE activity may not be fully constrained by genomic defences and shape the evolutionary trajectory of the species.

In this study, we screened whole-genome sequences of a global collection of 366 *P. nodorum* isolates for evidence of recent TE activity and genetic footprints of recent selection. TEs were exhaustivel y identified based on sequence similarity searches in three complete genomes of the species. We analyzed the TE load among populations in relationship with the demographic history of the pathogen. Finally, we identified TE loci potentially contributing to local adaptation and inferred the effectiveness of genomic defense mechanisms shaping TE variation.

## Material and Methods

### Populations characterized using Illumina whole-genome sequencing

We analyzed a total of 366 single spore isolates of *P. nodorum* sampled between 1991 and 2016. All isolates were collected from naturally infected wheat fields. The sampling locations included: Australia (2001 and 2010; *n* = 23), Brazil (year unknown; *n* = 1), Canada (2005; *n* = 1), Finland (year unknown; *n* = 1), Iran (2005 and 2010; *n* = 20), South Africa (1995; *n* = 23), Sweden (year unknown; *n* = 2), Switzerland (1999A and 1999B; *n* = 46), Arkansas (USA, 1995; *n* = 6), Georgia (USA, 2008; *n* = 5), Maryland (USA, 2008; *n* = 3), Minnesota (USA, 2002, 2003 and 2005; *n* = 14), New York (USA, 1991; *n* =21), North Carolina (USA, 2008; *n* = 9), North Dakota (USA, 1998, 2003, 2005, 2007, 2008, 2010, 2016; *n* = 82), Ohio (USA, 2003; *n* =16), Oklahoma (USA, 2016; *n* =17), Oregon (USA, 1993 and 2011; *n* =28), South Carolina (USA, 2008; *n* = 4), South Dakota (USA, 2016; *n* = 8), Tennessee (USA, 2008; *n* = 5), Texas (USA, 1992; *n* = 27) and Virginia (USA, 2008; *n* = 4). Earlier publications referred to the Switzerland 1999B population as being of Chinese origin (Sommerhalder et al. 2006; Stukenbrock, Banke & McDonald 2006; McDonald et al. 2012, 2013; Pereira et al. 2017). A more recent study corrected the population origin (Pereira, Croll, et al. 2020). Illumina whole-genome sequence data was generated for all 366 isolates with paired-end sequencing and a read length of 100-150 bp. Raw data was accessed from the NCBI Short Read Archive under BioProject ID numbers PRJNA606320, PRJNA398070 and PRJNA476481 (Syme et al. 2018; Richards et al. 2019; Pereira, Croll, et al. 2020).

### Genome alignment, variant calling and quality filtering

Raw reads were trimmed for remaining Illumina adaptors and read quality was assessed using Trimmomatic version 0.36 (Bolger et al. 2014) with the following parameters: illuminaclip = TruSeq3-PE.fa:2:30:10, leading = 10, trailing = 10, slidingwindow = 5:10, minlen = 50. Trimmed reads were aligned against the reference genome established for the isolate Sn2000 (Richards et al. 2017) using the short-read aligner Bowtie2 version 2.3.3 (Langmead & Salzberg 2012) with the –very-sensitive-local option. PCR duplicates were marked using the MarkDuplicates option in Picard tools version 2.17.2 (http://broadinstitute.github.io/picard). All sequence alignment (SAM) files were sorted and converted to binary (BAM) files using SAMtools version 1.2 (Li et al. 2009). Single nucleotide polymorphism (SNP) calling and variant filtration were performed using the Genome Analysis Toolkit (GATK) version 3.8-0 (McKenna et al. 2010). We used HaplotypeCaller on each alignment file individually with the --emit-ref-confidence GVCF and -ploidy 1 options. Joint variant calls were produced using GenotypeGVCFs with the flag -maxAltAlleles 2. Finally, SelectVariants and VariantFiltration were used for hard filtering SNPs failing the following cut-offs: QUAL > 200; QD > 10.0; MQ > 20.0; –2 < BaseQRankSum < 2; – 2 < MQRankSum < 2; –2 < ReadPosRankSum < 2. We kept only bi-allelic SNPs and with a maximum genotype missingness of 10% using vcftools version 0.1.15 (Danecek et al. 2011).

### Phylogenomic and population structure analyses

We inferred the evolutionary history of all 366 *P. nodorum* isolates using two phylogenetic tree construction methods. A maximum likelihood (ML) tree was estimated for all isolates using the GTRCAT model in RAxML version 8.2.12 (Stamatakis 2014). We performed 100 rapid bootstraps with 20 ML searches. The best ML tree was chosen based on support values. We analyzed evidence for reticulation among *P. nodorum* genotypes by building a Neighbor-Net (NN) network with SplitsTree4 version 4.15.1 (Huson & Bryant 2006). The ML tree and the NN network were visualized using the online tool iTOL version 4 (Letunic & Bork 2019). VCF files were converted to RAxML input (phylip) and SplitsTree4 input formats (nexus) using PGDSpider version 2.1.1.5 (Lischer & Excoffier 2012).

Population structure was inferred based on a principal component analysis (PCA) in TASSEL version 5.2.56 and a model-based clustering implemented in STRUCTURE version 2.3.4 (Pritchard et al. 2000; Bradbury et al. 2007). We performed the PCA based on SNP data filtered for a minor allele frequency above 5% and based on TE frequency insertion sites filtered for a minor allele frequency above 1%. We visualized the two first PCs using the *ggplot2* package in R (R Core Team 2019; Wickham 2009). For STRUCTURE analyses, we used an admixture model independent of prior population information and with correlated allele frequencies. The algorithm ran with a burn-in length of 50’000 and a simulation length of 100’000 Markov chain Monte Carlo (MCMC) repetitions. We explore a range of *K*between 1 and 10, with 10 repetitions per *K*. The most likely number of populations (*K*) was estimated using the Delta K (ΔK) method (Evanno et al. 2005) implemented in the R package *pophelper* version 2.3.0 (Francis 2017). For all phylogenetic and STRUCTURE analyses, we randomly selected SNPs at an average distance of 15 kb using the vcftools parameter --thin 15’000 to reduce linkage disequilibrium among loci and computational demands.

### Population genetics and selective sweeps

Allele frequencies and nucleotide diversity (p) per site were determined using the options --freq and --site-pi in vcftools (Danecek et al. 2011). Histograms for minor allele spectrum were generated in ggplot2. Linkage disequilibrium (LD) decay was estimated per population based on 50 kb windows on chromosome 1 using r^2^ with the option --hap-r2 in vcftools (Danecek et al. 2011). We performed Tajima’s D neutrality tests (Tajima 1989) using the vcftools --haploid flag and analyzed non-overlapping bins of 1 kb across the entire genome (Danecek et al. 2011). Selective sweeps were identified using a likelihood-based detection method implemented in the program SweeD version 3.0 using the option -folded (Pavlidis et al. 2013). We ran SweeD 3.0 individually per chromosome for grids of 1 kb. If LD decay was slow in a population (*i.e*. r^2^ ≥ 0.2 at 10 kb), we merged together adjacent genomic regions under selection if these were separated by less than 20 kb. We analyzed regions of interest by adding 10 kb on each side of the window identified by SweeD.

### Analysis of regions with signatures of selection using gene ontology

Genomic regions showing signatures of selection were used as coordinates in BEDtools version 2.29.0 (Quinlan & Hall 2010) to intersect annotated genes in the Sn2000 reference strain annotation (Richards et al. 2017). Genes were annotated for protein family (PFAM) domain and gene ontology (GO) terms using interproscan version 5.36-75.0 with default parameters and a local pre-calculated match lookup service (Jones et al. 2014). Protein secretion signals were predicted using SignalP version 4.01 (Nielsen 2017), Phobius (Käll et al. 2004) and TMHMM version 2.0 (Krogh et al. 2001). GO enrichment analyses were performed using the packages GSEABase version 1.35.0 and GOstats version 2.38.1 in R (Falcon & Gentleman 2007). We used the false discovery rate of 5% as a cut-off and minimum GO term size of 5 for the hypergeometric test.

### Transposable element consensus sequence identification and classification

To obtain consensus sequences for TE families, we performed individual runs of RepeatModeler version 1.0.8 on the three complete genomes Sn2000, Sn4 and Sn79-1087. Identified sequences were annotated based on GIRI Repbase using RepeatMasker version 4.0.7 (Richards et al. 2017; Bao et al. 2015; Smit et al. 2013). Further classification and processing of consensus sequences was performed using WICKERsoft (Breen et al. 2010). We screened the three complete genomes for copies of the above detected consensus sequences with BLASTN filtering for sequence identity of >80% over >80% of the length of the sequence (Altschul et al. 1997). We then added flanks of at least 300 bp both up- and downstream of each sequence. We made multiple sequence alignments using ClustalW and defined TE boundaries by visual inspection before updating consensus sequences (Higgins & Sharp 1988). If possible, consensus sequences were classified according to the presence and type of terminal repeats, superfamily-specific start and end bases or target site duplications, as well as homology of encoded proteins using BLASTX in the NCBI nr database. We excluded duplicated consensus sequences using dotter for inspection (Sonnhammer & Durbin 1995). We assigned consensus sequence names according to the three-letter naming system (Wicker et al. 2007). Two predicted TE families showed strong length polymorphism and only weak sequence similarity between all individual insertions. Using BLASTN on individual insertions, we found that the sequences matched one of three different regions of the entire consensus sequences. We visualized the consensus sequence alignment with genoPlotR version 0.8.9 in R (Guy et al. 2010) and used BLASTN to identify matching regions in the Sn2000, Sn4, and Sn79-1087 complete genomes (Richards et al. 2017; Morgulis et al. 2008; Zhang et al. 2000). To re-define consensus sequences, we proceeded as described above. In a second round of TE annotation, we focused on protein-coding sequences matching previously identified fungal TE superfamilies. We screened the three complete genomes for matches of protein sequences representative of each superfamily from other fungi using tBLASTN. We filtered hits for a minimal alignment length of 80 bp and a sequence similarity >25%. Identified sequences were retrieved including 300 bp flanking sequences. Hits were analyzed for their matching sequencing with dotter and grouped into families based on visual inspection. We made further multiple sequence alignments using ClustalW and defined TE boundaries by visual inspection. Finally, the TE family consensus sequences from the two methods above were used to annotate the reference genomes using RepeatMasker version 4.0.7 (Richards et al. 2017; Bao et al. 2015; Smit et al. 2013). We estimated the guanine-cytosine (GC) content as G+C/G+C+A+T on the TE families retained in the consensus sequence: the fasta consensus file was converted to EMBOSS sequence query using EMBOSS seqret (Madeira et al. 2019), and ran on EMBOSS geecee (https://www.bioinformatics.nl/cgi-bin/emboss/geecee). TE presence-absence annotation in the global collection of isolates was performed with the method proposed by Linheiro & Bergman (2012), with ngs_te_mapper version fb23590200666fe66f1c417c5d5934385cb77ab9 (https://github.com/bergmanlab/ngs_te_mapper/commit/fb23590200666fe66f1c417c5d5934385cb77ab9) implemented in R. Dependences to ngs_te_mapper pipeline were bwa version 0.7.17-r1188 (Li & Durbin 2009), to map Illumina reads, and samtools version 1.2 (Li et al. 2009), to perform output file conversion.

Depth of coverage can impact robust TE discovery, therefore we set a minimum depth of 10X, which allows recovery of ≥ 90% of identified TEs across populations (Supplementary Figure S1). As a consequence, we removed 18 isolates and retained a total of 348 isolates for downstream TE polymorphism analyses. We clustered nearby TE insertions into a single locus if the insertions (i) belonged to the same TE family and (ii) were located within 100 bp distance. Clustering of TE insertion sites was performed using the package GenomicRanges version 1.38.0 implemented in R (Lawrence et al. 2013).

## Results

### Global population structure and phylogenomics

We analyzed genome sequencing data for 366 *P. nodorum* isolates and identified a total of 487’477 high-confidence SNP markers with a minor allele frequency of 5%. We identified three main groups by performing principal component analyses (Figure 1A; Supplementary Figure S2). Isolates from Oklahoma (USA) formed a distinct group flanked by the two largest clusters. Unsupervised Bayesian clustering analyses revealed a similar pattern of high admixture among genotypes from different continents with the optimal number of clusters being *K* = 2 (Figure 1B; Supplementary Figure S3). At *K* = 3, isolates from Iran and the northern United States (North Dakota, South Dakota, and Minnesota) grouped as genetic cluster 2, while isolates from the southern United States shared membership with cluster 1. Genotypes from Australia and South Africa were assigned mainly to cluster 3. Genotypes from Swiss populations and Oklahoma showed significant admixture. We used genome-wide SNPs to infer a maximum-likelihood tree and identified four well-supported major clades largely independent of sampling origin (Figure 1C). We found evidence of reticulation indicative of recombination based on a SplitsTree network (Figure 1D). The network also showed that multiple populations harbored groups of highly similar genotypes indicative of recent ancestry. Both the maximum-likelihood tree and the SplitsTree network identified four major genetic groups including a group of isolates from Australia and South Africa, a group of isolates from Brazil, Finland, Sweden, Switzerland and the southern United States, as well as a group of isolates from Canada, Iran, and the northern United States. The fourth group is composed of isolates solely from Oklahoma (United States). Our results indicate significant levels of gene flow among populations from the same continent, as well as among a set of populations from different continents.

**Figure 1.**
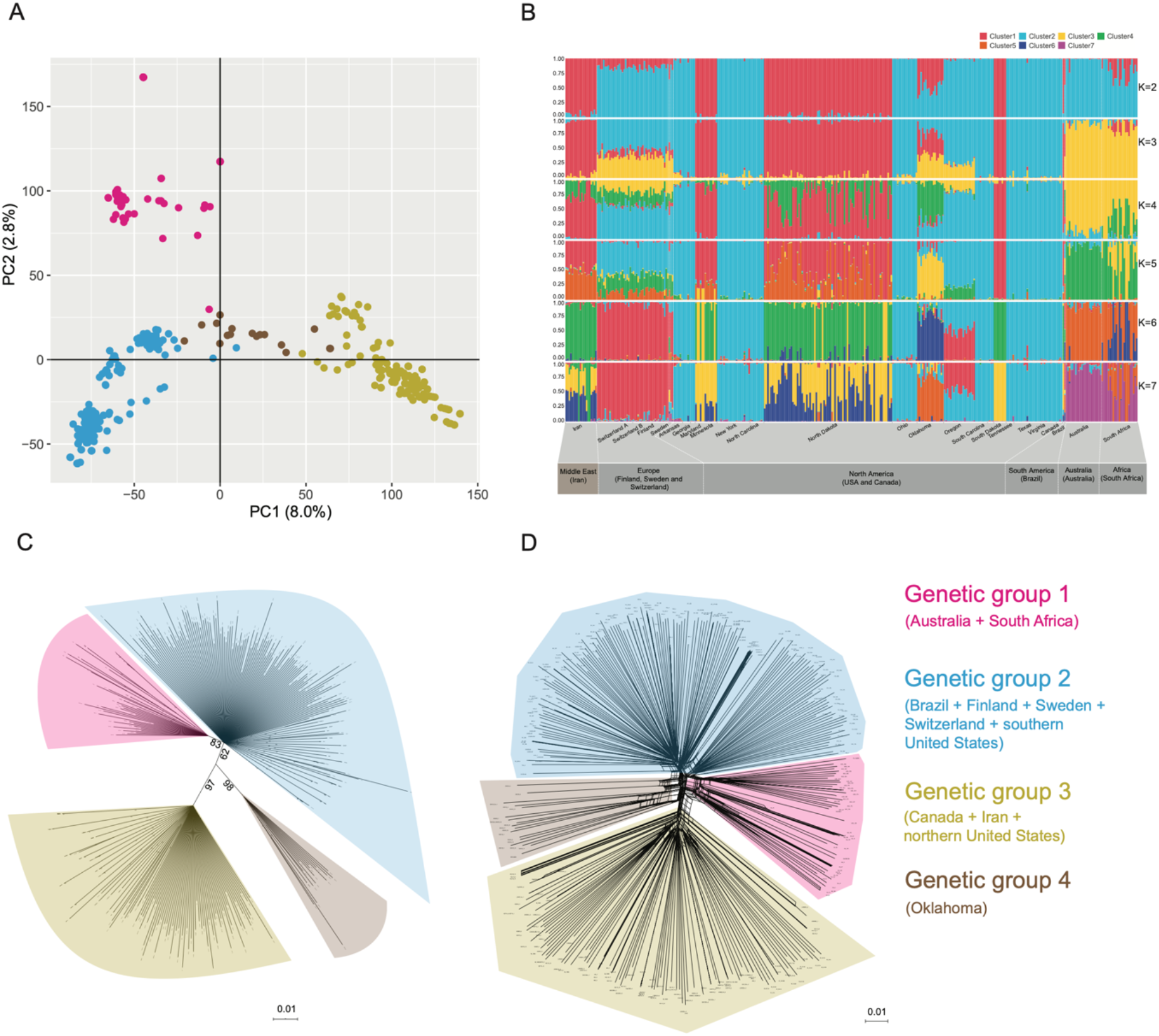
Global population structure and phylogenomics of *Parastagonospora nodorum*. (A) Principal component analysis (PCA) with dots representing individual isolates colored according to the genetic group. (B) Estimated population genetic structure. Each vertical bar represents one individual, colored according to cluster membership values. The cluster designation is represented by *K* numbers. Iran is highlighted as matching the region of origin of *P. nodorum*. (C) Maximum likelihood phylogenetic tree. (D) Phylogenetic network reconstruction. Population genetic structure and phylogenetic reconstructions were based on 2’278 genome-wide single nucleotide polymorphism (SNP) markers spaced evenly at 15 kb. The PCA was based on the complete SNP dataset.

### Population-level diversity and demography

We used nucleotide diversity and minor allele frequency spectra to quantify variation in genetic diversity across major genetic groups and local populations (Figure 2). Genetic group 3 (northern US and Iran) had the highest nucleotide diversity (4.64 X 10^−9^) but also the most considerable variation among sites (Figure 2A). North Dakota and Iran were highly diverse in contrast to South Dakota. Genetic groups 2 (mainly southern US) and 4 (Oklahoma) had intermediate levels of nucleotide diversity (4.19 X 10^−9^ and 4.02 X 10^−9^, respectively). Genetic group 1 had the lowest nucleotide diversity (3.84 X 10^−9^). We analyzed minor allele frequency spectra across populations to detect evidence for past demographic events (e.g., bottlenecks and expansions). Except for South Dakota, populations showed low-frequency alleles being more abundant than high-frequency alleles suggesting random mixing and low degrees of recent admixture (Figure 2B). Next, we estimated LD decay and found that the distance for r^2^ < 0.2 varied between ~2.2 kb in Switzerland 1999B and Australia to 44.1 kb in Oklahoma (Figure 2B). Most populations (10 out of 16) showed a fast LD decay (r^2^ < 0.2 within 10 kb), indicating high levels of diversity and ongoing recombination. The slow LD decay in South Africa and Oklahoma may indicate recent admixture or more dominant roles of asexual reproduction.

**Figure 2.**
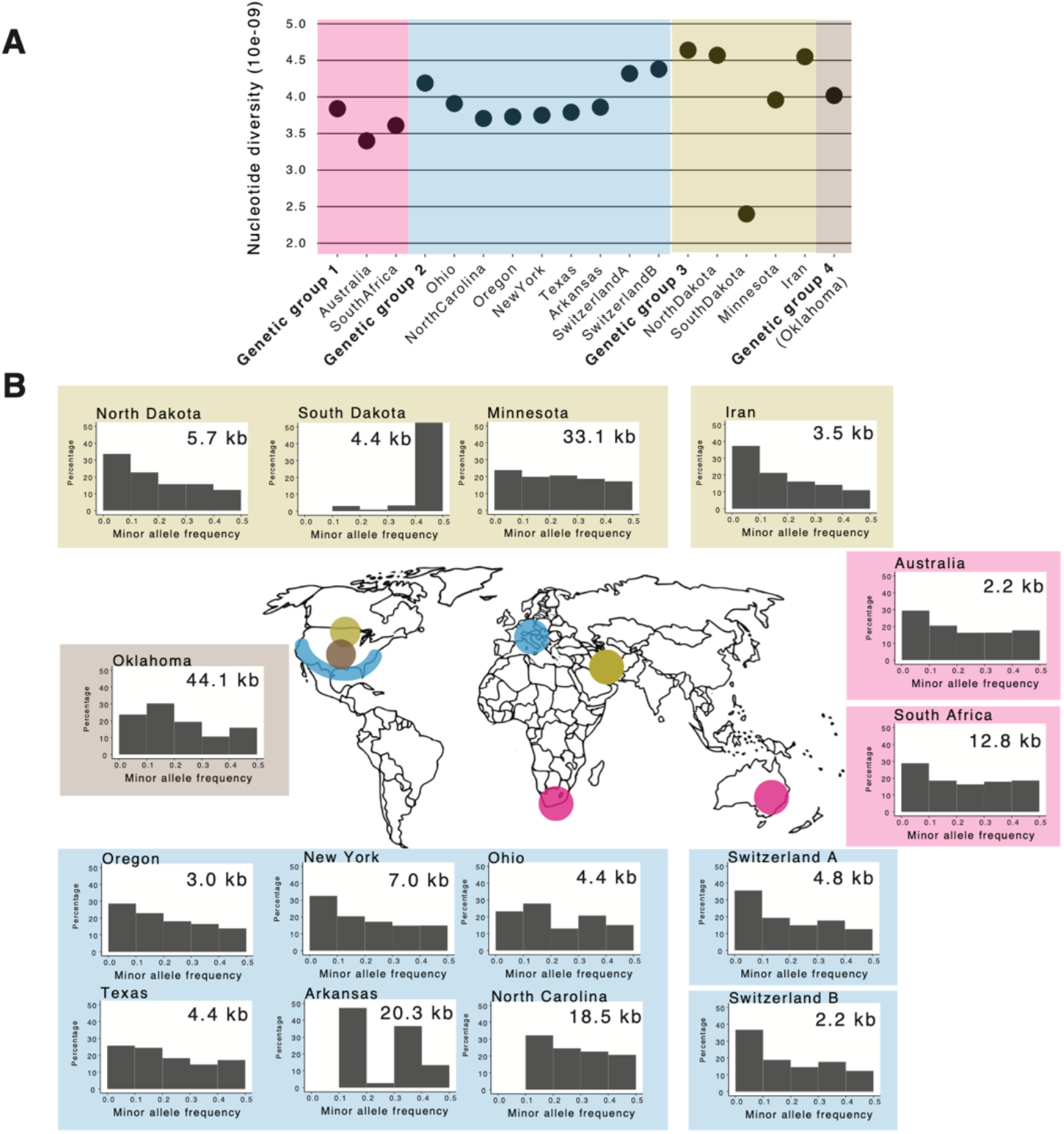
Worldwide analyses of demography and population diversity. (A) Nucleotide diversity for genetic groups and sampling locations. Colors represent different genetic groups. (B) Minor allele frequency spectrum for individual sampling locations. Sampling locations are represented on the map, highlighting northern United States and Iran in yellow; Oklahoma in brown; southern United States and Switzerland in blue, and Australia and South Africa in pink. The number in the top right corner of histograms represents the distance for linkage disequilibrium to decay below *r*^2^ < 0.2. All estimates are based on SNPs on chromosome 1.

### TE landscape and insertion dynamics among populations

Adaptation to local conditions may emerge from genetic variation produced by TE mediated sequence rearrangements. Here, we performed *de novo* TE annotation using three complete genomes of *P. nodorum* (Richards et al. 2017) and combined this information into a single set of TE consensus sequences. Our consensus annotation identified 25 TE families in *P. nodorum* and revealed that the genomes Sn2000, Sn4, and Sn79-1087 are composed of 4.40%, 4.23% and 1.57% TEs, respectively (Figure 3; Supplementary Table S1). The TE density among the three reference genomes was heterogeneous (Figure 3A). The reference isolate Sn79-1087 showed an overrepresentation of TEs mostly in subtelomeric regions. TEs in Sn2000 and Sn4 showed TE blocks along chromosomes with a particularly high density on chromosome 10 (Figure 3A). Among the TE classes, class I elements comprised 3.76%, 3.37%, and 0.82% of the genome and class II elements comprised 0.50%, 0.72%, and 0.42% of the genome in Sn2000, Sn4 and Sn79-1087, respectively. These findings are consistent with the previous estimates (Hane et al. 2007; Richards et al. 2017). Moreover, we show that complete genomes vary in the content and density of TEs highlighting the interested to screen TE dynamics at the population level.

**Figure 3.**
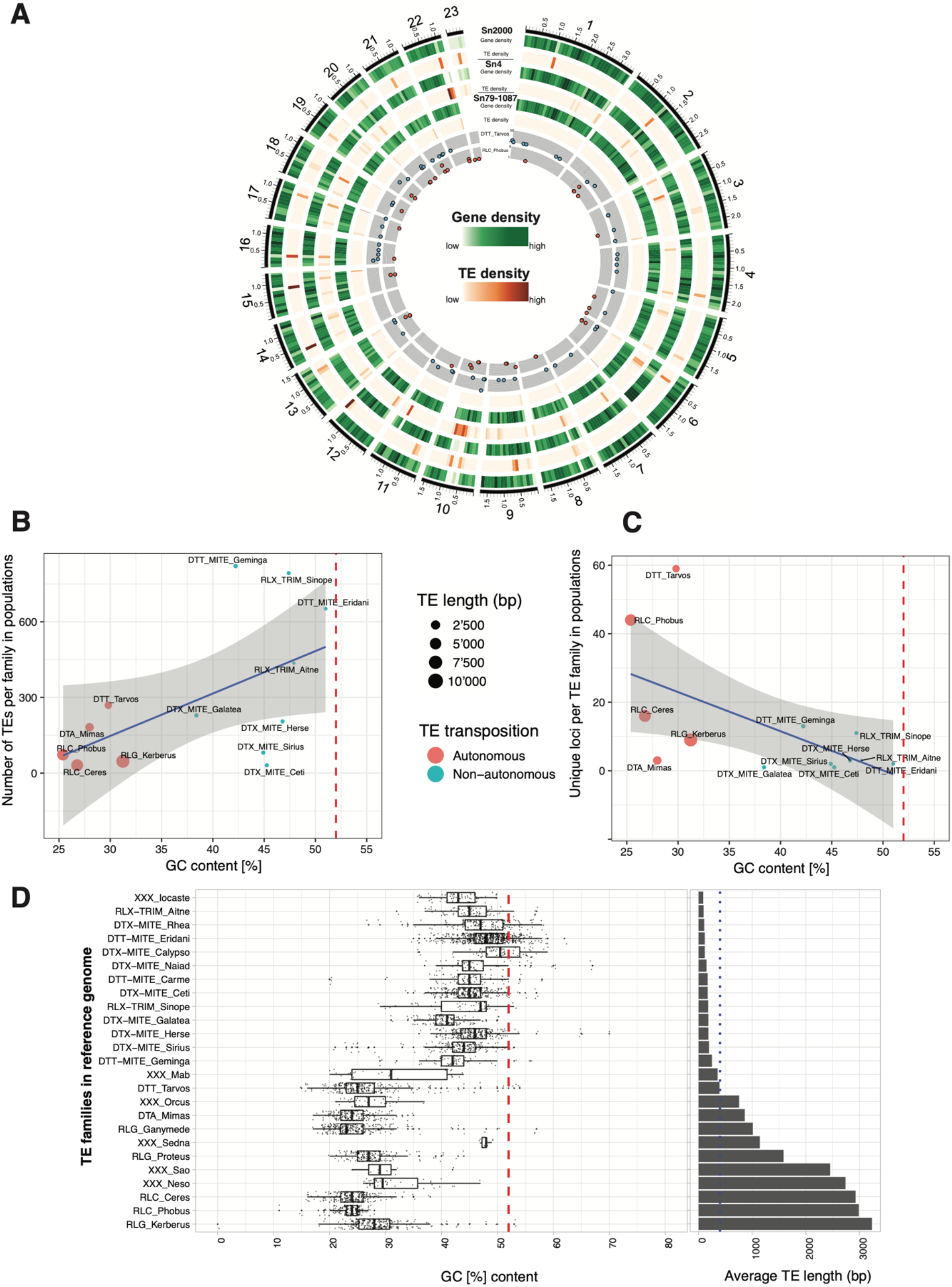
Genome-wide detection of transposable elements (TEs). (A) Annotation of the completely assembled reference genomes. The outer black ring delimitates the 23 chromosomes and positions are indicated in Mb. The gene density is calculated for 100 kb windows (darker green for higher densities). The TE density was calculated for 100 kb windows (darker orange for higher densities). Inner grey rings highlight the occurrence of TE families with the highest number of loci in the genome (DTT_Tarvos with blue circles) and the highest degree of singleton (RLC_Phobus with red circles).The position on the y-axis shows the relative difference in the allele frequencies among populations. (B) Linear correlation plot of GC content and TE copy number across populations. (C) Linear correlation plot of distinct loci per TE family across populations. (D) Analyses of TE copies in the three reference genomes for GC content and the average TE length. Red vertical dashed lines indicate the genome-wide average GC content (52%) and the dotted blue vertical line shows the 400 bp threshold.

We inferred population-level variation in TE activity by analyzing the 348 isolates with genome sequences for evidence of newly inserted or deleted TEs. We identified 3’850 TE insertions across all isolates with the insertions clustered into 167 unique loci in the genome (Supplementary Figure S4A). Recently inserted TEs were mainly DNA transposons (class II; *n*=2’470) and included fewer retrotransposons (class I; *n*=1’380). At the order level, we found only terminal inverted repeats (TIR) and long terminal repeats (LTR), respectively (Figure 4A). Superfamilies were composed mainly of elements classified as class II *Tc1-Mariner* (*n*=1’742), followed by class II *hAT* (*n*=182) and class I *Copia* (*n*=104; Figure 4A). The remaining 1’229 LTRs and 546 TIRs were not conserved enough to assign a superfamily. We found a stark difference between autonomous (*n*=602) and non-autonomous elements (*n*=3’248; Figure 4A; Supplementary Figure S4B). Non-autonomous elements included 2’019 miniature inverted repeats (MITE; 62.1 %) and 1’229 terminal repeat retrotransposons in miniature (TRIM; 37.9%). The total count of recent TE insertions in a population represents the load generated through TE activity. TEs were highly homogeneous between populations and genetic groups, with the highest average numbers of new insertions in isolates of genetic group 1 (South Africa and Australia; mean = 13.1 insertions) and the lowest in genetic group 4 (Oklahoma; mean = 8.5 insertions; Figure 4C; Supplementary Figure S5). The TE superfamily composition is highly similar among genetic groups with a predominance in *Tc1-Mariner* and an unknown LTR (Figure 4D). Recently inserted *Gypsy* elements were found only in genetic groups 2 and 3. At the family level, insertions of DTX_MITE_Sirius were mostly found at a single locus on chromosome 19 (99%), and insertions of DTA-Mimas were mostly found at a single locus on chromosome 23 (67%). Chromosome 23 was identified as an accessory chromosome in *P. nodorum* (Ohm et al. 2012; Richards et al. 2017). Similarly, TE families DTX_MITE_Ceti and DTX_MITE_Galatea were found only at a single locus on chromosomes 12 and 19, respectively. Most variation in TE loci numbers was driven by DTT_Tarvos (269 copies in total among isolates distributed across 59 loci, Figure 3A). The RLC_Phobus showed a high degree of singletons with different 44 loci (Figure 3A). Finally, TE copy expansion was driven by non-autonomous elements DTT_MITE_Geminga (821 copies), RLX_TRIM_Sinope (793 copies), and DTT_MITE_Eridani (652 copies). The species-wide screens of recently inserted TEs show that populations harbor fairly balanced TE loads with only a minor divergence in TE composition.

**Figure 4.**
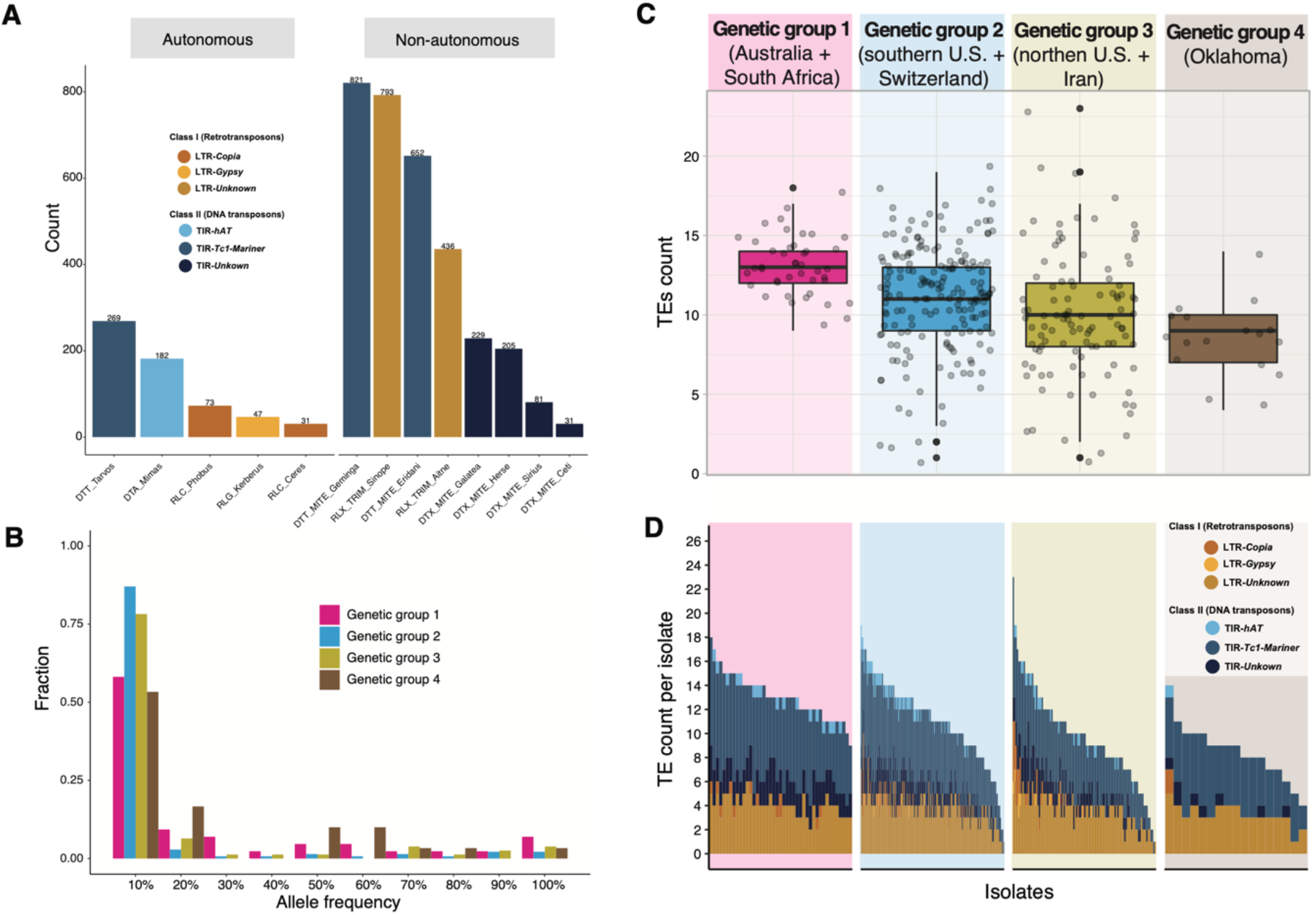
Classification and differential load of transposable elements (TE) among genetic groups. (A) Barplots of TE counts according to family and transposition mode. (B) TE insertion frequency distribution per genetic group. (C) The average number of TEs per isolate within genetic groups. (D) TE counts per isolate and TE family across genetic groups. Each bar represents a single isolate and colors identify TE families.

### Differential activity among TEs and evidence for purifying selection

*P. nodorum* has an active genome defence called RIP, that is inducing C:G to T:A mutations in recently duplicated genomic sequences. The preferential AT-mutation bias of RIP will gradually lower the GC content in RIP-affected genomic regions (Watters et al. 1999; Hane & Oliver 2008). We estimated GC content as a proxy for RIP activity at the level of TE families. Active TEs not affected by RIP are expected to have a GC content comparable to the genome-wide average of coding sequences. We considered the population-level TE activity as the total number of copies per TE family across the 348 isolates. Interestingly, we found that TE families with higher copy numbers across the populations have higher GC content than low-copy TE families (Figure 3B). The three most expanded TE families (DTT_MITE_Geminga, RLX_TRIM_Sinope, and DTT_MITE_Eridani) showed the highest GC content (mean GC of 41.8, 43.7 and 48.2%, respectively). In contrast, TE families with copies divided across more loci were lower in GC content, while those in fewer unique loci showed higher GC content (Figure 3C). The majority of TEs with high GC content in both analyses were non-autonomous MITEs and TRIMs. We also found consistent variation in GC content within TE families, with TEs of short length usually being of higher GC content (e.g., MITEs and TRIMs) (Figure 3D). Overall, recently active TEs in the *P. nodorum* genome are mostly of short length and constituted of non-autonomous elements.

TE frequencies across the genome are impacted by the joint actions of TE activity, selection on the insertion, and demography. We analyzed whether the TE insertion frequency spectrum reflected neutral processes such as drift and migration, or selection. We performed a principal component analysis using TE presence at insertion loci as a genetic marker. In contrast to genome-wide SNPs, we found no geographical signal for TE insertion loci (Supplementary Figure S6A-B). The majority of all TEs were present at <10% frequency in the global set of isolates, and about 48.5% of all TEs were singletons (Supplementary Figure S7). We found a particularly pronounced shift towards singletons in genetic group 2, suggesting strong purifying selection acting against the TEs in this group (Figure 4B; Supplementary Figure S8).

### Genomic signatures of recent selective sweeps

We analyzed genetic groups for footprints of recent selective sweeps. We calculated the composite likelihood ratio (CLR) implemented in the software SweeD for genetic groups 1-3 separately (group 4 composed of Oklahoma was omitted given the high admixture signatures). We detected a total of 46 genomic regions with significant signatures of recent sweeps across all groups (Supplementary Figure S9; Supplementary Table S2). Selective sweeps were detected on all chromosomes except for chromosomes 1, 14, and 16. The size of the regions showing signatures of selective sweeps ranged from 20-40.7 kb. Interestingly, no overlaps in selective sweep regions were found among the genetic groups 1-3, indicating strong heterogeneity in selection pressures and local adaptation. In genetic group 1, a total of 16 genomic regions with sweep signatures were detected on chromosomes 2, 4-6, 8, 9, 11, 12, 15, 18-20, and 23 based on a 99.9% outlier threshold (Figure 5A, Supplementary Table S2). The average length of the sweep regions was 21.3 kb spanning a total of 341.5 kb or 0.9% of the genome. The selective sweep region with the highest likelihood score (136) was located on chromosome 2 (Figure 5B). We analyzed Tajima’s D in the same region and found a positive but below-average value for the chromosome (Figure 5C). Linkage disequilibrium decay was slower in the sweep region with an average of r^2^=0.48 over ~20.8 kb compared to a decay of r^2^ < 0.2 within 5.7 kb for genetic group 1 (Figure 5E). The above-average linkage disequilibrium is consistent with a recent selective sweep. In genetic groups 2 and 3, we detected a total of 16 genomic regions with signatures of selection in total (Supplementary Figure S9; Supplementary Table S2).

**Figure 5.**
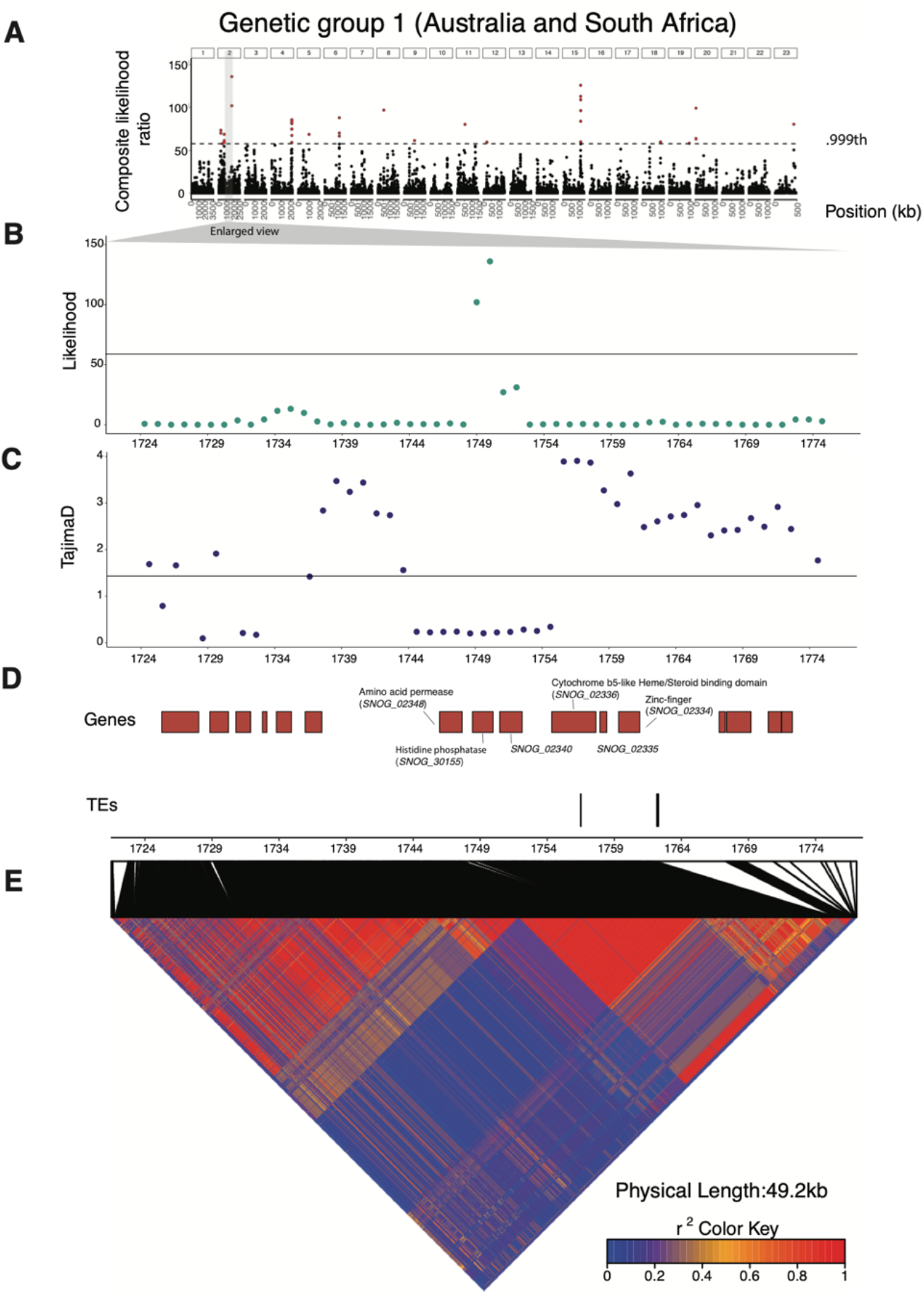
Selective sweep loci in genetic group 1. (A) Distribution of the composite likelihood scores (CLS) in windows of 1 kb. The dashed horizontal line indicates the 99.9^th^ percentile threshold and the yellow highlight shows the strongest sweep region. (B) Distribution of the CLS within the strongest sweep region. (C) Tajima’s *D* calculated for 1 kb windows. (D) Schematic of genes highlighting predicted function and transposable elements. (E) Heatmap of pairwise linkage disequilibrium *r*^2^ within the sweep region.

### Genes potentially underlying local adaptation

Globally, we found 334 genes in selective sweep regions (genetic groups 1-3; Supplementary Table S2). The number of genes per sweep region ranged from 2-22. In genetic group 1 (South Africa and Australia), we identified a total of 99 genes (Supplementary Table S3). The average number of genes per sweep region was 6.2 ranging from 3-10 genes. We tested whether genes in sweep regions were overrepresented for genes encoding specific protein functions. We found 27 significantly overrepresented GO terms, including basal metabolic processes (e.g., cellular macromolecule biosynthetic/metabolic process), nutrient mobilization (e.g., nitrate assimilation), and transcriptional regulation (e.g., regulation of transcription, zinc ion binding; Supplementary Table S4). The genomic region with the highest likelihood score in the genetic group 1 contained two genes with roles in transcription, including *SNOG_02334* (a Zinc-finger transcription factor) and *SNOG_30155* (a histidine phosphatase superfamily). We also identified genes encoding transporters, including *SNOG_02348* (amino acid permease) and *SNOG_02336* (cytochrome b5-like heme; Figure 5D). Overall, genes in sweep regions were found to control metabolic functions and regulatory processes.

Among the 99 genes, two genes (*SNOG_07292, SNOG_30828*) were previously ranked as strong candidate effectors (Syme et al. 2018). The genetic signal for a sweep found on chromosome 8 (from 520242 to 540242 bp) was in close proximity to the locus containing the necrotrophic effector *ToxA*. A detailed analysis revealed a strong LD with the downstream region of *ToxA* and several TE insertion loci (Supplementary Figure S10). The presence of *Copia, Gypsy, and Tc1-Mariner-like* elements were previously reported at the *ToxA* locus (McDonald et al. 2019), here we identified additional MITEs and unclassified TEs (Supplementary Figure S10). We examined whether the 99 genes located in selective sweep regions from genetic group 1 were in close physical proximity of TEs, but we detected no over-representation of TEs in proximity of these genes compared to the genomic background (Supplementary Figure S11). Our findings show that the recent insertion dynamics of TEs in *P. nodorum* populations are unlikely to have driven recent selective sweeps. However, the insertion of individual TEs in proximity to known effector genes may well have had an impact on the evolution of virulence.

## Discussion

The TEs activity can be a major source of genetic variation in the fungal genome. Yet, we lack a comprehensive view how evolutionary forces including genetic drift and selection act on TE dynamics in natural populations. Here, we analyzed the intra-species population genetic diversity, genome-wide dynamics of TEs and signatures of selection in a worldwide collection of the plant pathogen *P. nodorum*. This species exhibits relatively low population subdivisions consistent with recent gene flow among continents. The TE compositions and copy numbers were very similar among populations. Despite strong evidence for highly active genomic defences (Hane & Oliver 2008, 2010), we identified recent TE activity in the species pool. TEs do not appear to be the main drivers of recent adaptive evolution and are rather under purifying selection. Yet, we show that some high-copy TEs have sequence signatures consistent with an escape from genomic defence mechanisms.

For this work, we significantly expanded previous population genomic analyses of *P. nodorum* for both geographic coverage and sample size (Richards et al. 2019; Pereira, McDonald, et al. 2020; Pereira, Croll, et al. 2020). Our results corroborate evidence for genetic admixture among North America, Europe (Switzerland), and the Middle East (Iran) consistent with earlier analyses based on microsatellite loci (Stukenbrock, Banke & McDonald 2006). High levels of gene flow among geographically widespread pathogen populations can lead to maladaptation and counteract local adaptation (Croll & McDonald 2017). Yet, gene flow can be particularly advantageous for plant pathogens in agroecosystem because agricultural practices and key susceptibility genes can be shared globally among crop growing areas (Croll & McDonald 2017; Richards et al. 2019). We indeed found high genetic diversity within and among field *P. nodorum* populations, with a rapid LD decay reflecting frequent sexual recombination. The slow LD decay found in more recently founded populations (e.g., South Africa) most likely reflects recent admixture. The historic expansion of wheat cultivation across the globe was likely accompanied by *P. nodorum* spreading from the Fertile Crescent to most continents. However, the global expansion of agriculture and pathogens likely imposed strong genetic bottlenecks in newly founded populations (McDonald & Linde 2002; Oliver et al. 2012). Besides a reduction in genetic diversity, such demographic effects can influence genome-wide TE activity in some species (Stritt et al. 2018; Oggenfuss et al. 2020). We found no striking differences in TE abundance among populations. The genetic group 1, comprising the more recently founded populations of Australia and South Africa, showed similar TE composition as genetic group 2 (including Iranian isolates). Populations of the wheat pathogen *Z. tritici* showed striking expansion in most TE superfamilies (Stukenbrock, Banke, Javan-Nikkhah, et al. 2006; Oggenfuss et al. 2020). As *Z. tritici* and *P. nodorum* are thought to share common domestication origins and subsequent dispersal routes, differences in how TEs were impacted may be due to differences in genomic defenses (e.g. cytosine DNA methylation polymorphism in *Z. tritici* (Möller et al. 2020). Plant populations can similarly be shaped by bottleneck effects, but TEs did not show pronounced activation as found for example in the wild grass *Brachypodium distachyon* (Stritt et al. 2018). TE activity shaped by demography highlights how genome-wide TE proliferation is complex and can be governed by stochastic effects.

We identified differences in the genome-wide TE distribution among the three completely assembled and re-annotated *P. nodorum* reference genomes. Compartmentalization of repetitive regions in the genome was previously shown (Richards et al. 2017), and can mediate local adaptation by causing gene gains and losses (Richards et al. 2019). However, we found little evidence for TEs playing a major role on selective sweeps leading to local adaptation or sweep regions enriched in virulence factors in our populations. Nevertheless, the low amount of repetitive sequences and extensive signatures of RIP strongly suggest that the *P. nodorum* genome is largely devoid of TE activity (Hane et al. 2007, 2007; Hane & Oliver 2008). This is consistent with the view that TE multiplication in a genome is determined by the balance between the TE’s transposition potential and the effectiveness of genomic defenses (Muszewska et al. 2017). Here, we show that *P. nodorum* TE families exhibit a striking variation in GC content, which is positively correlated with the TE copy numbers in the genome. The correlation suggests that an increase in TE copy numbers is associated with weak RIP activity on these TEs. RIP guards the genome against TE proliferation by targeting duplicated DNA (*e.g*., recent copies of TEs) but the RIP intensity can vary among TE families (Santana et al. 2014; Amselem et al. 2015).

In *N. crassa*, the action of RIP is impaired in duplicated sequences shorter than ~400 bp (Yeadon & Catcheside 1995; Watters et al. 1999), and with an identity below 80% (Cambareri et al. 1991). We found that high-copy TE families with high GC content are mostly MITEs and TRIMs. These elements generally lack coding sequences (Wicker et al. 2007) and were on average <400 bp in *P. nodorum*. Hence, the MITEs and TRIMs may have escaped RIP through their compactness. Both groups of TEs were found to mediate genome evolution and modulate gene expression across eukaryotes (Witte et al. 2001; Santiago et al. 2002; Naito et al. 2009; Fouché et al. 2020). In addition, MITEs are thought to increase in copy numbers despite genomic defenses with possible benefits to DNA transposons (Feschotte et al. 2003; Feschotte & Pritham 2007). MITEs and TRIMs are ubiquitous in the pangenome of the wheat pathogen *Z. tritici*, which has a similar genome size but more active TEs (Badet et al. 2020). MITEs in *Z. tritici* become de-repressed upon encountering stressful conditions such as during the colonization of the host plant (Fouché et al. 2020). The MITE activity in *P. nodorum* could also be a consequence of posttranscriptional regulation. In rice, short interfering RNA was found to preferentially suppress MITEs expression (Nolan 2005; Lu et al. 2012). Genomic regions showing signatures of recent selective sweeps did not contain recently inserted TEs. Whether the activity of MITEs and TRIMs in *P. nodorum* is contingent on stressful conditions and whether transposition activity creates adaptive variation remains unknown.

In this study, we resolved the global population structure and retraced TE activity in the repressive genome of an important wheat pathogen. TEs can play a major role in plant pathogen adaptation by generating adaptive genetic variation to resist pesticides or to overcome host defence mechanisms. Key pathogenicity genes show tightly associated expression patterns with TEs in several plant pathogens. However, the selfish nature of TEs can also impose severe costs on the pathogen. Population genetic analyses informed by highly contiguous genome sequences are powerful tools to disentangle the evolutionary equilibrium between TE activation and repression.

## Acknowledgments

We thank Cécile Lorrain for comments and suggestions on a previous version of the manuscript. The Genetic Diversity Center (GDC) of ETH Zurich and the Functional Genomics Center in Zurich provided sequencing facilities. This study was financed in part by the Coordenação de Aperfeiçoamento de Pessoal de Nível Superior – Brasil (CAPES) – Finance Code 001.

