## Supplementary Information for "The population genomics of transposable element activation in the highly repressive genome of an agricultural pathogen"

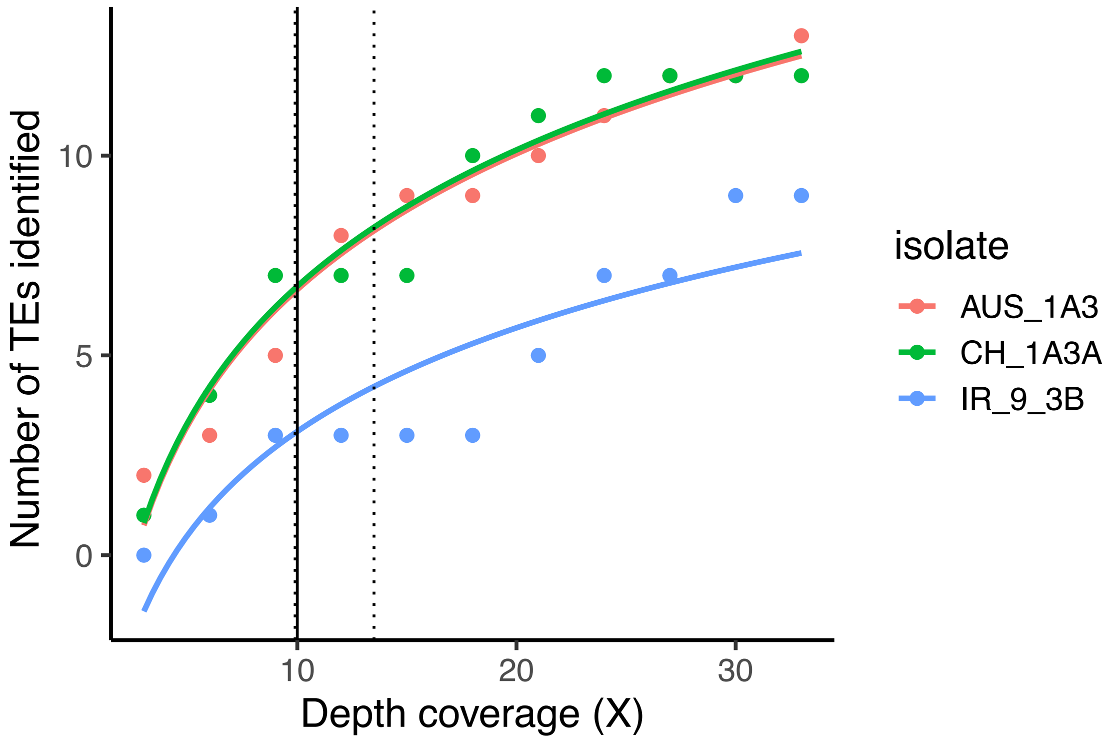

**Supplementary Figure 1. Coverage down-sampling to assess transposable element (TE) discovery.** Vertical dashed lines indicate a 90% interval for the detection of the maximum number of TEs. The solid black line indicates the minimum threshold set at 10X.

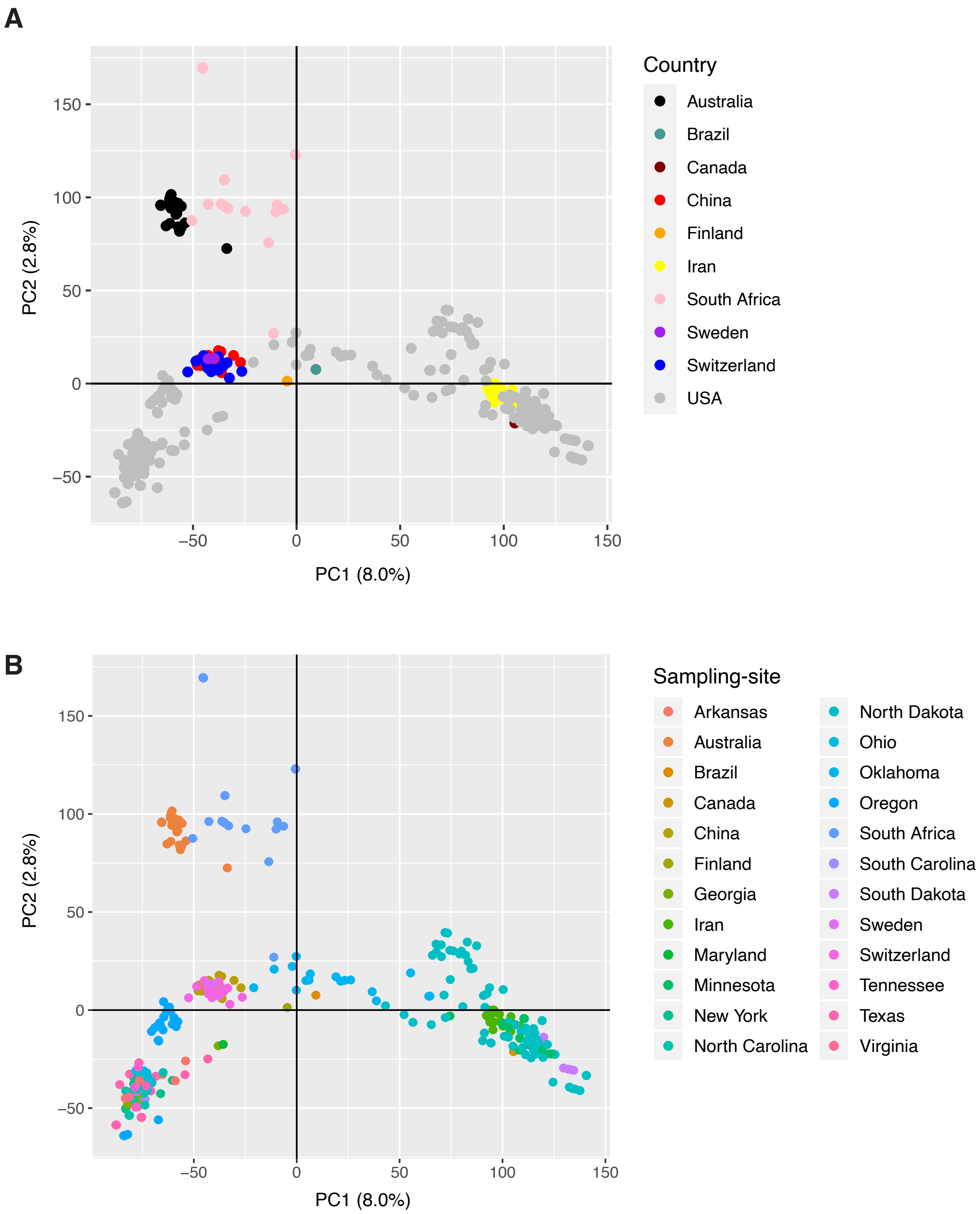

**Supplementary Figure 2.** Principal component analysis (PCA) based on the complete SNP dataset. (A) PCA with dots representing the country of sampling. (B) PCA with dots representing the sampling sites.

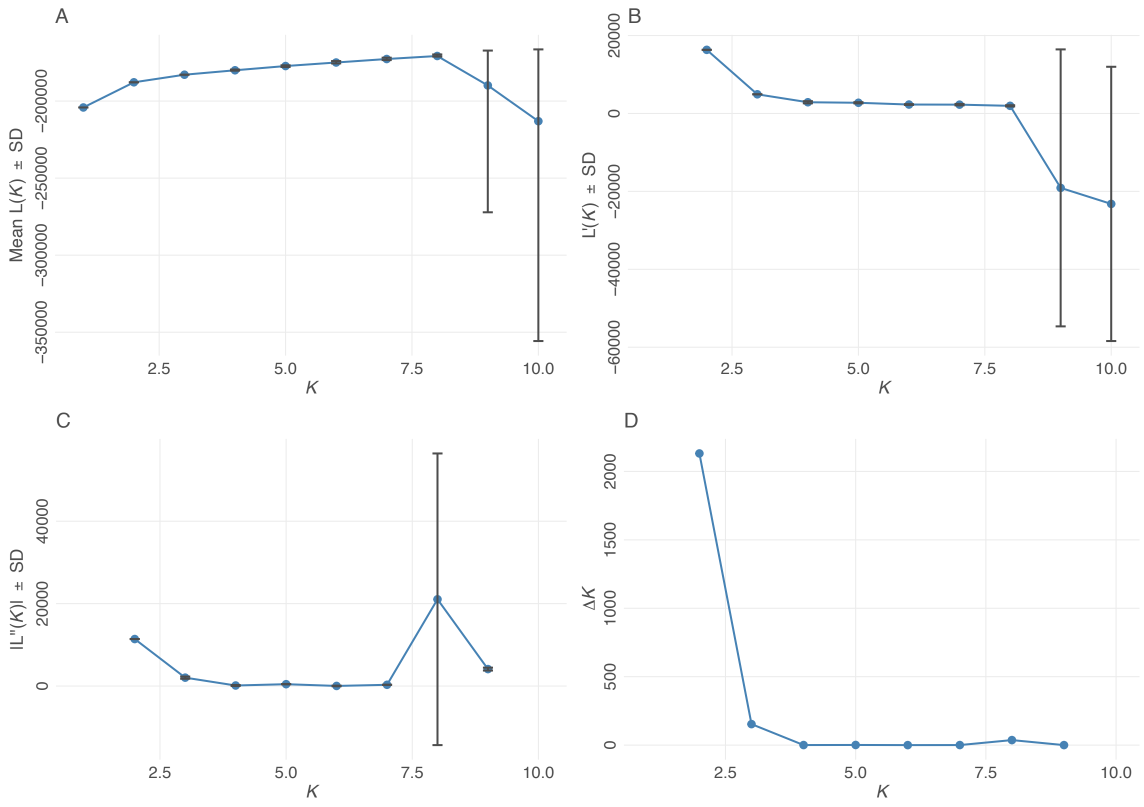

**Supplementary Figure 3. Cluster assignment statistics using STRUCTURE.** (A) Average likelihood and variance for each *K* value. (B) The ratio of the likelihood distribution. (C) The modular value of the second-order rate of change for the likelihood trend. (D) Mean delta *K* plot varying between *K*=2 and *K*=9.

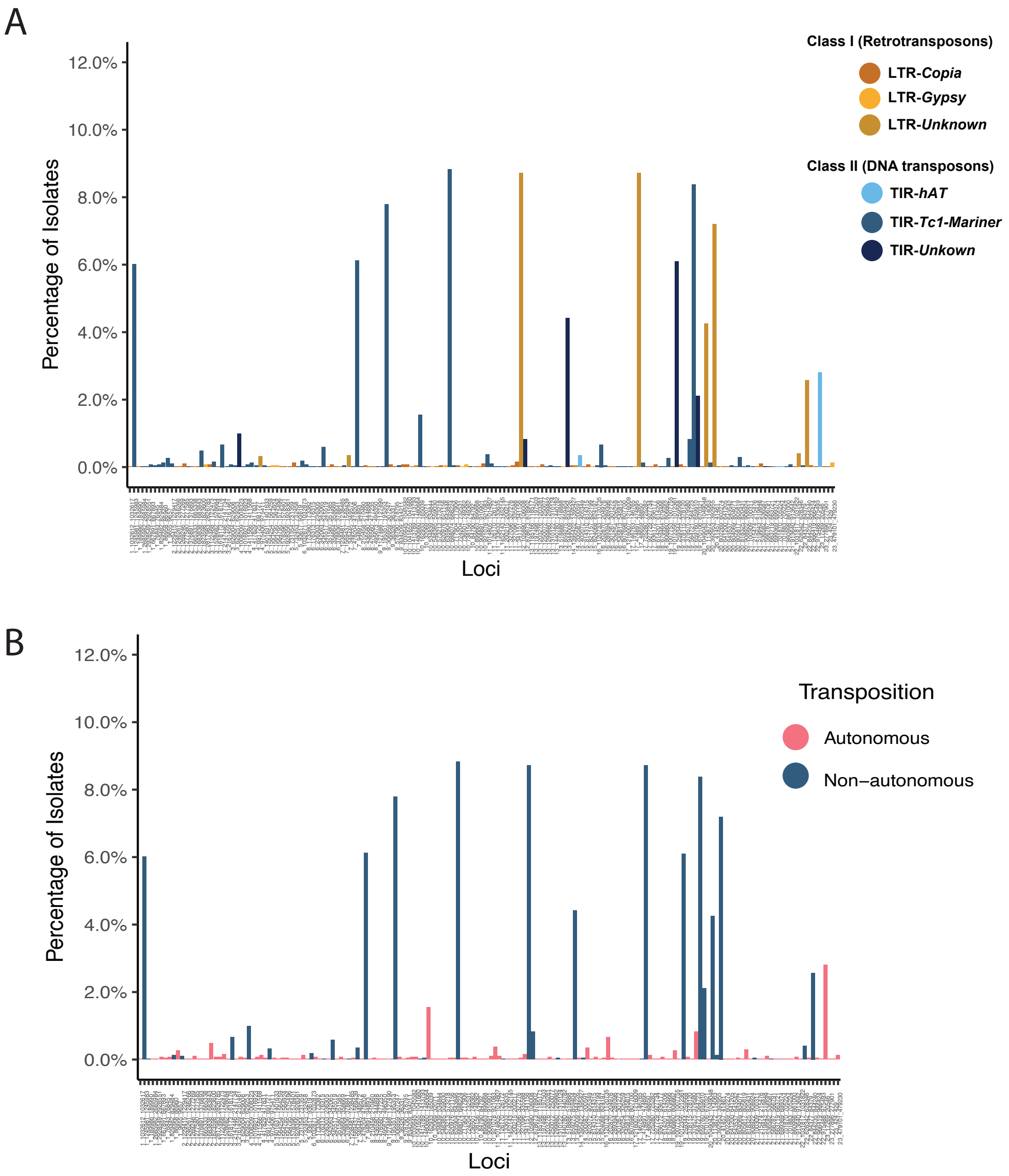

**Supplementary Figure 4. Transposable element (TE) frequencies across insertion loci in the genome.** Colors represent (A) family classification and (B) transposition mode.

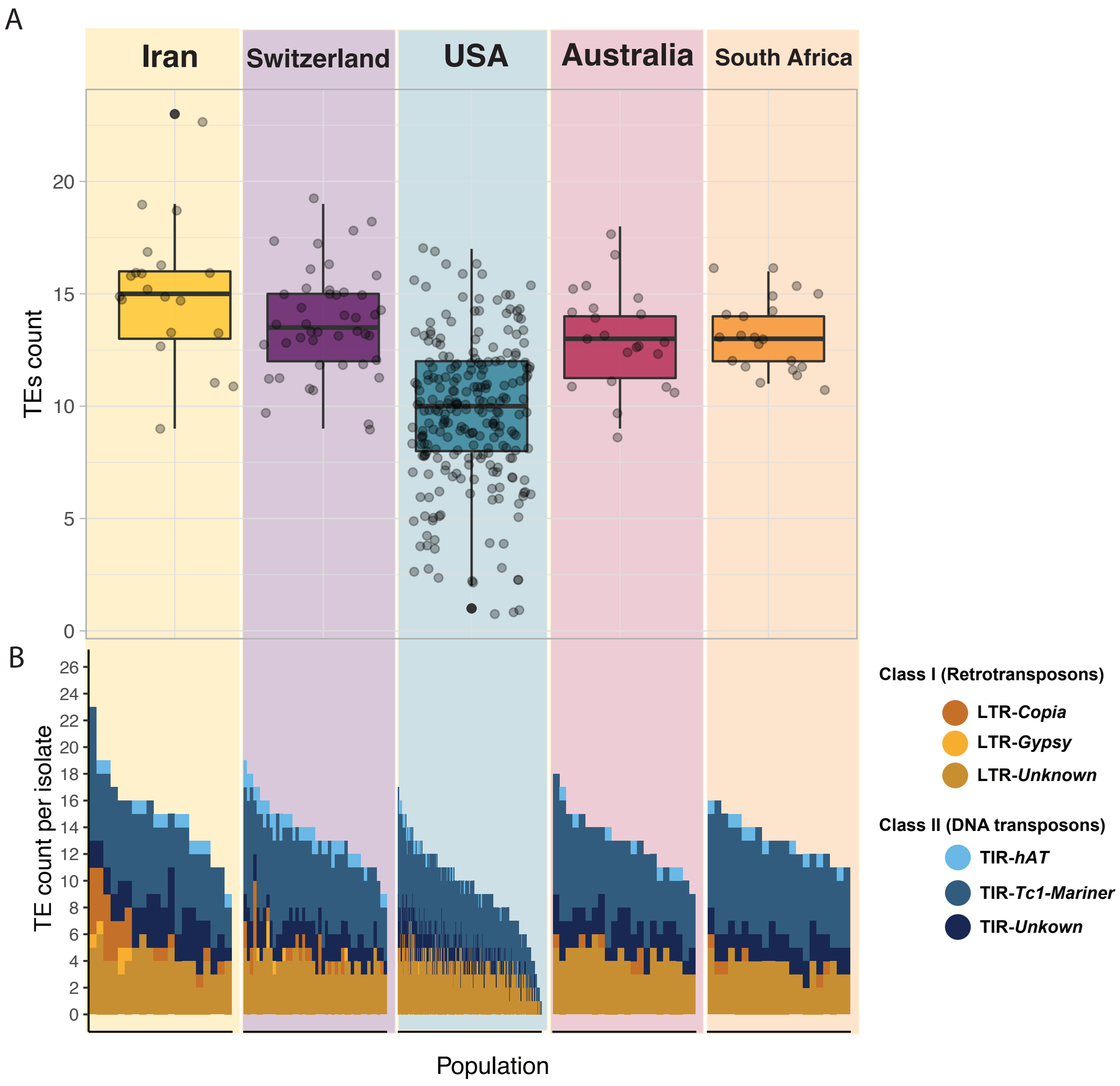

**Supplementary Figure 5. Classification and differential load of transposable elements (TE) among populations.** (A) The average number of TEs per isolates within populations. (B) Count of TEs per isolate and TE family across populations.

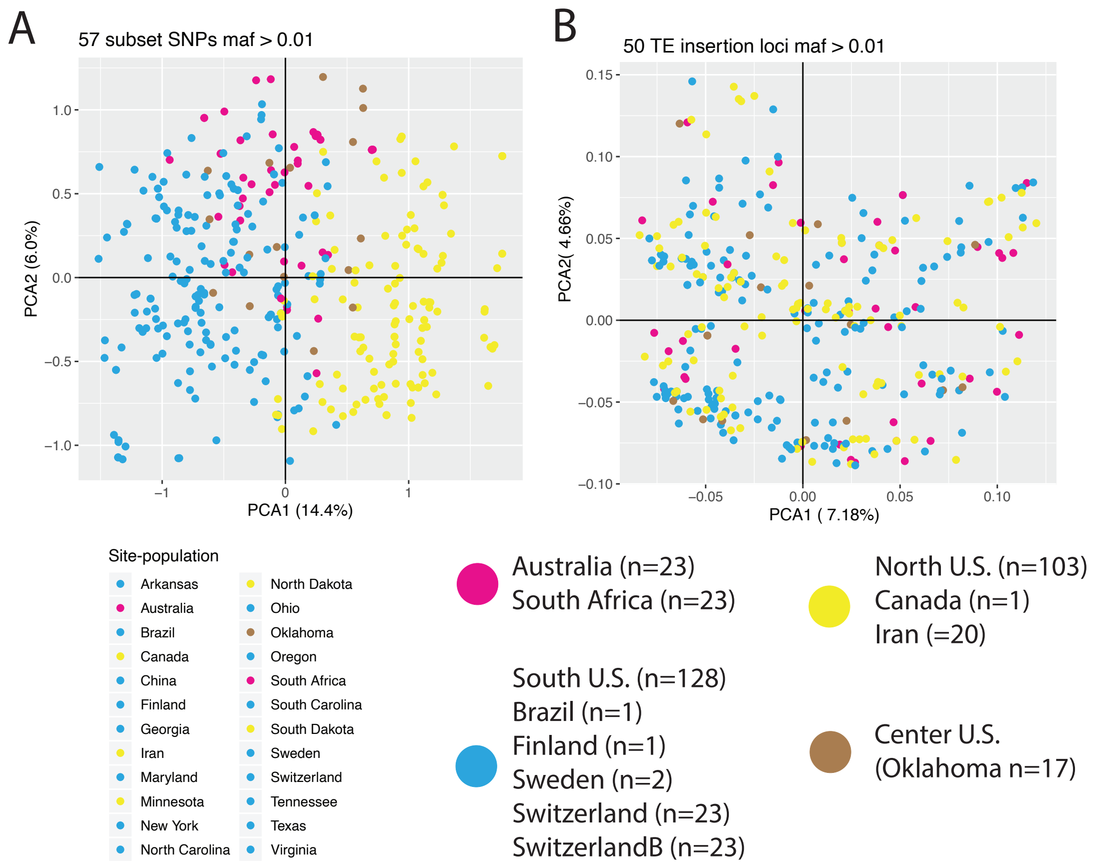

**Supplementary Figure 6. Population inferences based on genome-wide single nucleotide polymorphisms (SNP) and transposable element (TE) presence/absence genotypes at different loci.** The number of loci were intentionally set to be comparable between the two marker types. (A) Principal component analysis (PCA) based on 57 randomly selected SNP markers. (B) PCA based on 50 TE insertion loci. The different dots represent different isolates and colors identify the four major genetic groups.

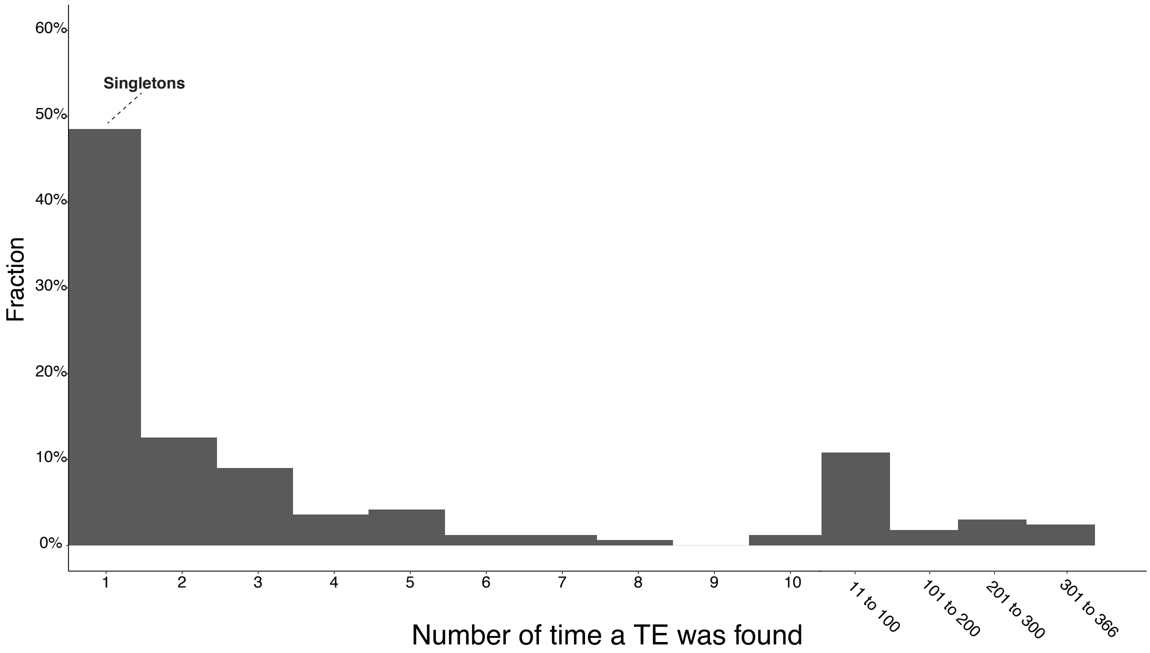

**Supplementary Figure 7. Transposable element (TE) presence frequencies at insertion loci across all isolates.** Singletons are highlighted and identify loci where only a single isolate carried a TE among all isolates.

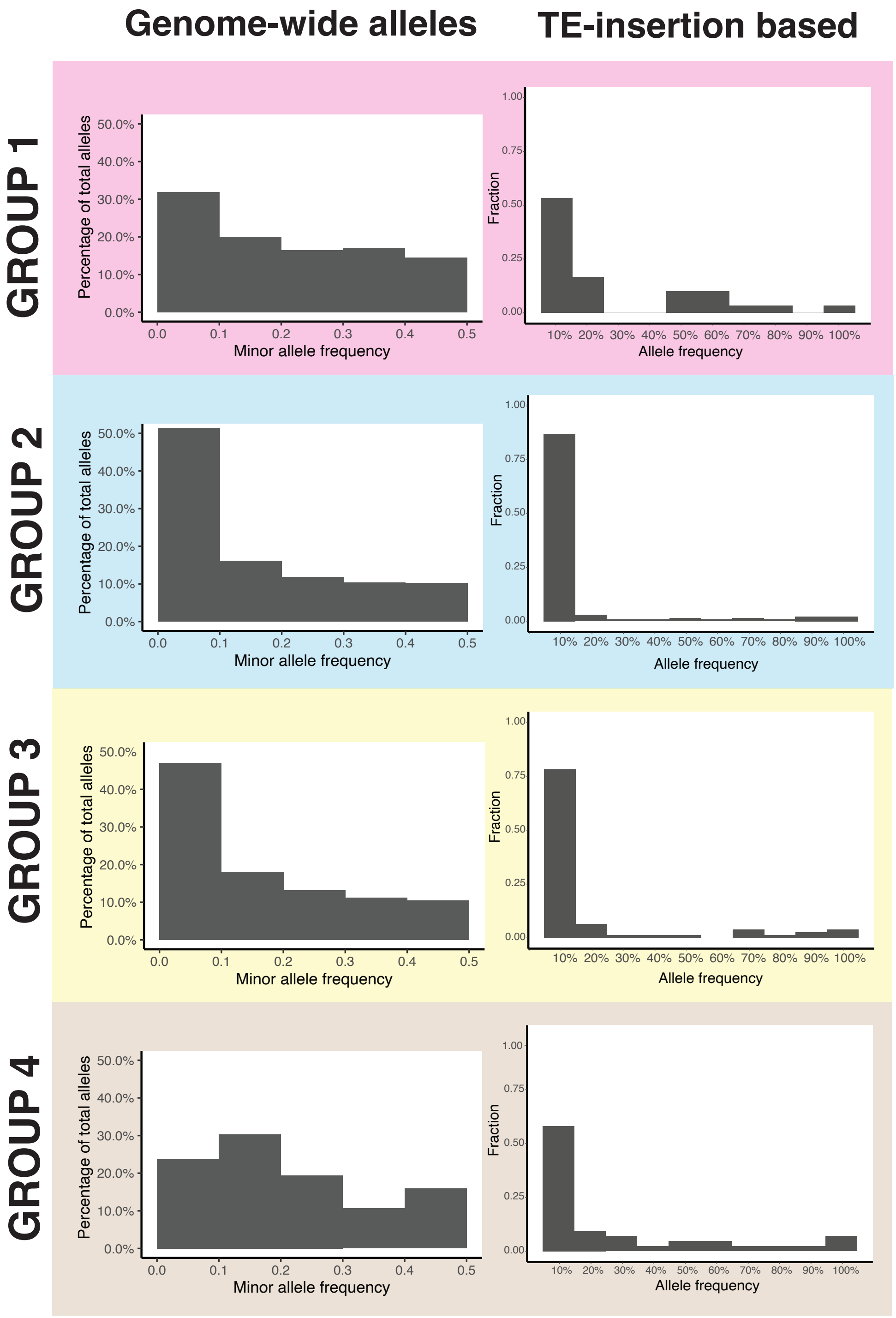

**Supplementary Figure 8. Minor allele frequency distribution from genome-wide single nucleotide polymorphisms (SNP) (left column) and transposable element insertion frequencies (right column) across the four genetic groups (rows).**

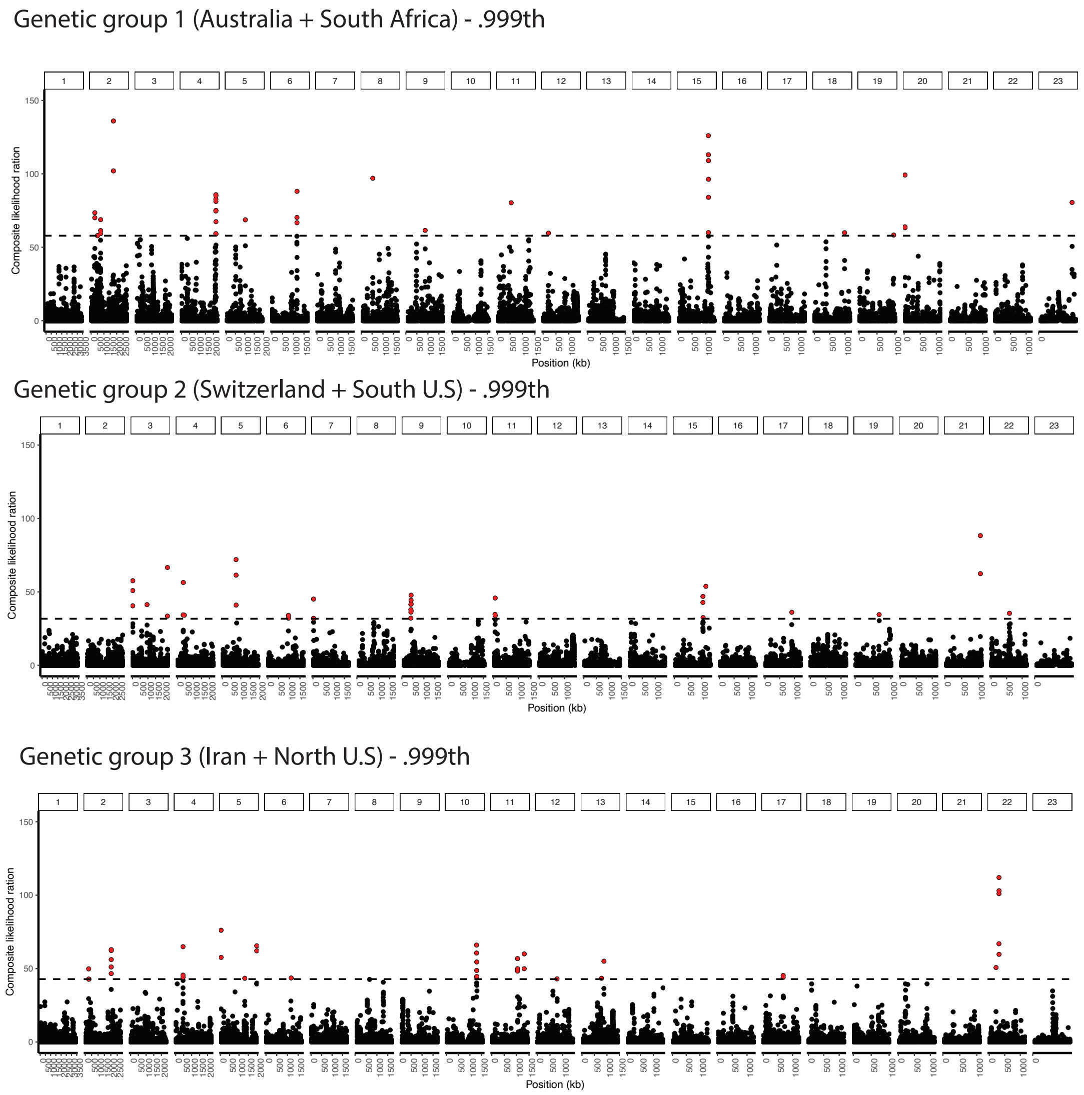

**Supplementary Figure 9. Differences in selective sweep regions among genetic groups of *Parastagonospora nodorum.*** Horizontal dashed lines represent the 99.9$^{th}$ percentile threshold of significance for each group. Loci above the threshold are highlighted in red.

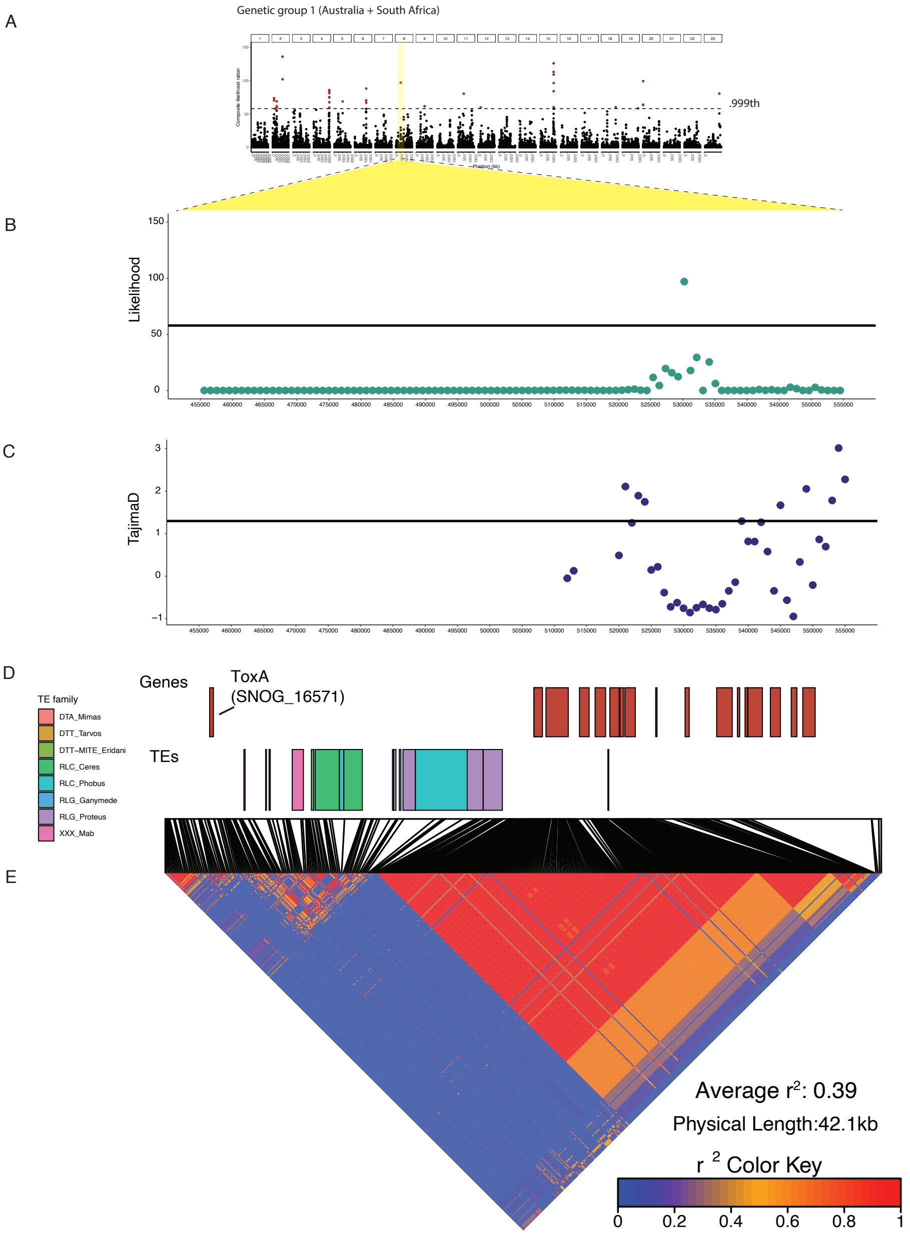

**Supplementary Figure 10. Genomic analyses of a selective sweep region on chromosome 8 in genetic group 1.** (A) Composite likelihood scores (CLS) in windows of 1 kb. The dashed horizontal line indicates the 99.9$^{th}$ percentile threshold and yellow highlights the strongest sweep region. (B) CLS within the strongest sweep region. (C) Tajima's *D* calculated for 1 kb windows. (D) Diagram of predicted genes and function. Annotated transposable elements colored by families. (E) Heatmap of pairwise linkage disequilibrium $r^2$ within the sweep region.

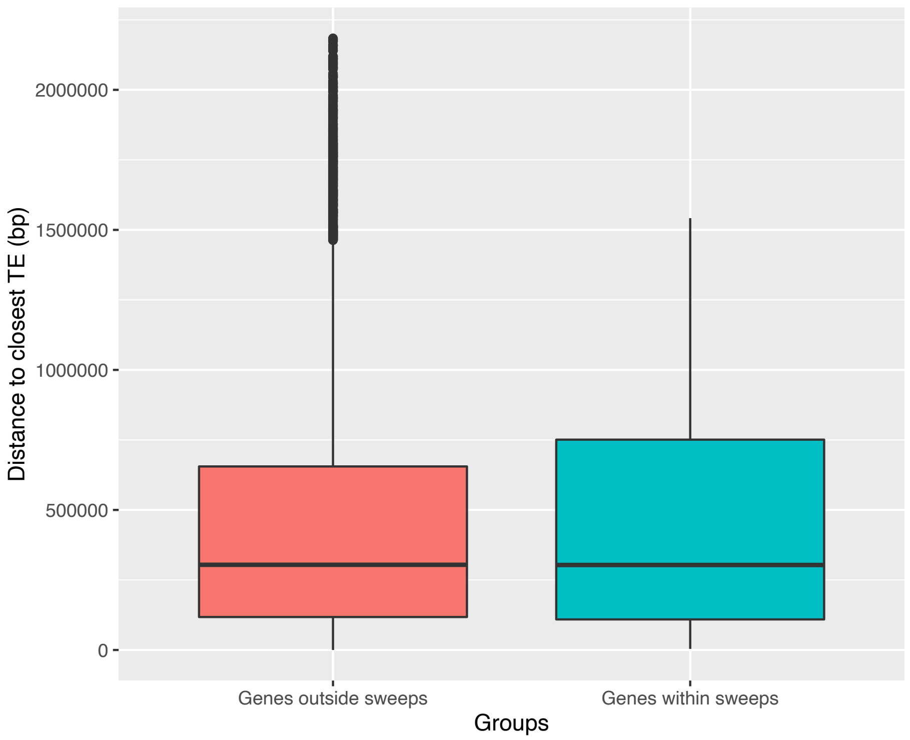

**Supplementary Figure 11. The distance of genes to the closest transposable elements.** Krustal-Wallis rank-sum test; *p*-value = 0.311.

**Supplementary Table 1.** RepeatMasker summary of transposable elements (TEs) detected in the three complete *Parastagonospora nodorum* reference genomes (Sn2000, Sn4 and Sn79-1087).

| TE Classification[1] | | | | Sn2000 genome length: 37154224 bp | | Sn4 genome length: 37695105 bp | | Sn79-1087 genome length: 34781891 bp | |
|---|---|---|---|---|---|---|---|---|---|
| TE_class | TE_order | TE_superfamily | TE_family | length(bp) | percentage | length(bp) | percentage | length(bp) | percentage |
| I | LTR | *Copia* | RLC_Ceres | 268696 | 0.72 | 477913 | 1.27 | 10200 | 0.03 |
| | | | RLC_Phobus | 405527 | 1.09 | 181568 | 0.48 | 35094 | 0.10 |
| | | Gypsy | RLG_Ganymede | 71353 | 0.19 | 79213 | 0.21 | 150642 | 0.43 |
| | | | RLG_Kerberus | 523458 | 1.41 | 410040 | 1.09 | 16378 | 0.05 |
| | | | RLG_Proteus | 120402 | 0.32 | 116363 | 0.31 | 65464 | 0.19 |
| | | unknown | RLX-TRIM_Aitne | 3085 | 0.01 | 2514 | 0.01 | 2972 | 0.01 |
| | | | RLX-TRIM_Sinope | 3456 | 0.01 | 3051 | 0.01 | 5096 | 0.01 |
| II | TIR | *hAT* | DTA_Mimas | 42375 | 0.11 | 109314 | 0.29 | 13512 | 0.04 |
| | | Tc1-Mariner | DTT_Tarvos | 33657 | 0.09 | 54585 | 0.14 | 22205 | 0.06 |
| | | | DTT-MITE_Carme | 5523 | 0.01 | 5050 | 0.01 | 4470 | 0.01 |
| | | | DTT-MITE_Eridani | 26278 | 0.07 | 26349 | 0.07 | 26767 | 0.08 |
| | | | DTT-MITE_Geminga | 6015 | 0.02 | 5014 | 0.01 | 5077 | 0.01 |
| | | unknown | DTX-MITE_Calypso | 2846 | 0.01 | 2698 | 0.01 | 2791 | 0.01 |
| | | | DTX-MITE_Ceti | 17014 | 0.05 | 18049 | 0.05 | 17796 | 0.05 |
| | | | DTX-MITE_Galatea | 8876 | 0.02 | 8383 | 0.02 | 8800 | 0.03 |
| | | | DTX-MITE_Herse | 19072 | 0.05 | 18568 | 0.05 | 19127 | 0.05 |
| | | | DTX-MITE_Naiad | 3569 | 0.01 | 3286 | 0.01 | 3278 | 0.01 |
| | | | DTX-MITE_Rhea | 6147 | 0.02 | 6250 | 0.02 | 6699 | 0.02 |
| | | | DTX-MITE_Sirius | 14172 | 0.04 | 14825 | 0.04 | 14096 | 0.04 |
| unknown | unknown | unknown | XXX_Iocaste | 1980 | 0.01 | 2012 | 0.01 | 1929 | 0.01 |
| | | | XXX_Mab | 5586 | 0.02 | 6090 | 0.02 | 1789 | 0.01 |
| | | | XXX_Neso | 30568 | 0.08 | 26933 | 0.07 | 73174 | 0.21 |
| | | | XXX_Orcus | 3852 | 0.01 | 5064 | 0.01 | 21998 | 0.06 |
| | | | XXX_Sao | 9606 | 0.03 | 1693 | 0.00 | 15671 | 0.05 |
| | | | XXX_Sedna | 1713 | 0.00 | 7991 | 0.02 | 1716 | 0.00 |
| | | | Total | 238849 | 0.64 | 322154 | 0.85 | 260895 | 0.75 |

[1] TE_class (I = retrotransposons, II = DNA transposon); TE_order (LTR = long terminal repeat, TIR = terminal inverted repeat).

**Supplementary Table 2.** List of selective sweep regions and number of overlapping genes found in each of the three genetic groups.

| Genetic group | Chromosome | start (bp) | end (bp) | length (bp) | Number of genes |
|---|---|---|---|---|---|
| 1 | 2 | 280224 | 301212 | 20988 | 6 |
| 1 | 2 | 489663 | 510651 | 20988 | 10 |
| 1 | 2 | 734668 | 756644 | 21976 | 3 |
| 1 | 2 | 1739385 | 1760373 | 20988 | 6 |
| 1 | 4 | 2141207 | 2169014 | 27807 | 4 |
| 1 | 5 | 1017824 | 1037824 | 20000 | 8 |
| 1 | 6 | 1162062 | 1184015 | 21953 | 4 |
| 1 | 8 | 520242 | 540242 | 20000 | 8 |
| 1 | 9 | 866643 | 886643 | 20000 | 6 |
| 1 | 11 | 535736 | 555736 | 20000 | 7 |
| 1 | 12 | 200453 | 220453 | 20000 | 8 |
| 1 | 15 | 1085013 | 1109854 | 24841 | 5 |
| 1 | 18 | 975976 | 995976 | 20000 | 5 |
| 1 | 19 | 954019 | 974019 | 20000 | 8 |
| 1 | 20 | 16221 | 38169 | 21948 | 8 |
| 1 | 23 | 409903 | 429903 | 20000 | 3 |
| | total | | | 341489 | 99 |
| 2 | 3 | 8741 | 30731 | 21990 | 8 |
| 2 | 3 | 934055 | 954055 | 20000 | 4 |
| 2 | 3 | 2255364 | 2278349 | 22985 | 10 |
| 2 | 4 | 347622 | 372500 | 24878 | 3 |
| 2 | 4 | 412016 | 432016 | 20000 | 8 |
| 2 | 5 | 731552 | 756437 | 24885 | 6 |
| 2 | 6 | 951110 | 972087 | 20977 | 11 |
| 2 | 7 | 17594 | 44446 | 26852 | 5 |
| 2 | 9 | 398784 | 431121 | 32337 | 7 |
| 2 | 11 | 45623 | 73356 | 27733 | 11 |
| 2 | 15 | 1031730 | 1058509 | 26779 | 6 |
| 2 | 15 | 1139220 | 1159220 | 20000 | 8 |
| 2 | 17 | 901746 | 921746 | 20000 | 5 |
| 2 | 19 | 660562 | 680562 | 20000 | 10 |
| 2 | 21 | 1039649 | 1060592 | 20943 | 4 |
| 2 | 22 | 581112 | 601112 | 20000 | 8 |
| | total | | | 370359 | 114 |
| 3 | 2 | 257502 | 298248 | 40746 | 2 |
| 3 | 2 | 2022918 | 2047858 | 24940 | 2 |
| 3 | 4 | 448155 | 472058 | 23903 | 4 |
| 3 | 5 | 30489 | 51467 | 20978 | 5 |
| 3 | 5 | 1301733 | 1321733 | 20000 | 5 |

| | | | | | |
|---|---|---|---|---|---|
| 3 | 5 | 1943222 | 1964200 | 20978 | 5 |
| 3 | 6 | 1166544 | 1186544 | 20000 | 6 |
| 3 | 10 | 1276040 | 1304776 | 28736 | 10 |
| 3 | 11 | 1026821 | 1048754 | 21933 | 11 |
| 3 | 11 | 1296525 | 1317491 | 20966 | 11 |
| 3 | 12 | 763851 | 783851 | 20000 | 12 |
| 3 | 13 | 774301 | 794301 | 20000 | 13 |
| 3 | 13 | 871174 | 891174 | 20000 | 13 |
| 3 | 17 | 671566 | 692558 | 20992 | 17 |
| 3 | 22 | 208765 | 228765 | 20000 | 22 |
| 3 | 22 | 302105 | 325084 | 22979 | 22 |
| total | | | | 367151 | 121 |

**Supplementary Table 3.** List of genes found within selective sweeps in the genetic group 1. Chr = chromosome.

| Sweep ID | Chr | Gene | start (bp) | end (bp) | strand | Pfam | GOterms |
|---|---|---|---|---|---|---|---|
| Chr.2.1739385.1760373 | 2 | SNOG_02334 | 1760002 | 1760373 | - | Zinc-finger of mitochondrial splicing suppresso | |
| Chr.2.1739385.1760373 | 2 | SNOG_02335 | 1758608 | 1759120 | + | | |
| Chr.2.1739385.1760373 | 2 | SNOG_02336 | 1755006 | 1758307 | + | Cytochrome b5-like Heme/Steroid binding don Oxidoreductase NAD-binding domain, Mo-co o | GO:0042128\|GO:0016491\|GO:0055114\|GO:0016491\|GO:0030: |
| Chr.2.1739385.1760373 | 2 | SNOG_02340 | 1751112 | 1752794 | + | | |
| Chr.2.1739385.1760373 | 2 | SNOG_02348 | 1746625 | 1748308 | - | Amino acid permease | GO:0016020\|GO:0022857\|GO:0055085 |
| Chr.2.1739385.1760373 | 2 | SNOG_30155 | 1749077 | 1750626 | + | Histidine phosphatase superfamily (branch 2) | GO:0016791 |
| Chr.2.280224.301212 | 2 | SNOG_08013 | 280870 | 281769 | - | | |
| Chr.2.280224.301212 | 2 | SNOG_08016 | 286472 | 287671 | - | Ubiquitin carboxyl-terminal hydrolase | GO:0006511\|GO:0036459\|GO:0016579\|GO:0036459 |
| Chr.2.280224.301212 | 2 | SNOG_08017 | 287851 | 289075 | + | | GO:0016021 |
| Chr.2.280224.301212 | 2 | SNOG_08018 | 289756 | 290674 | - | NmrA-like family | |
| Chr.2.280224.301212 | 2 | SNOG_08019 | 291399 | 292640 | + | Fungal Zn(2)-Cys(6) binuclear cluster domain | GO:0000981\|GO:0005634\|GO:0006355\|GO:0008270 |
| Chr.2.280224.301212 | 2 | SNOG_08020 | 292777 | 293658 | - | | |
| Chr.2.280224.301212 | 2 | SNOG_08021 | 295297 | 296016 | - | | |
| Chr.2.280224.301212 | 2 | SNOG_08022 | 296679 | 297923 | + | Diacylglycerol kinase catalytic domain | GO:0016301\|GO:0003951 |
| Chr.2.280224.301212 | 2 | SNOG_08023 | 298143 | 299333 | - | | |
| Chr.2.280224.301212 | 2 | SNOG_08024 | 300600 | 301212 | + | Zinc finger protein | |
| Chr.2.489663.510651 | 2 | SNOG_08122 | 490187 | 491706 | + | Phosphotransferase enzyme family | |
| Chr.2.489663.510651 | 2 | SNOG_08128 | 502036 | 503394 | + | Protein kinase domain | GO:0004672\|GO:0005524\|GO:0006468\|GO:0005524\|GO:0004( |
| Chr.2.489663.510651 | 2 | SNOG_08130 | 508863 | 510651 | - | Fungal specific transcription factor domain | GO:0003677\|GO:0005634\|GO:0006351\|GO:0008270\|GO:0000! 0008270 |
| Chr.2.734668.756644 | 2 | SNOG_08238 | 734669 | 735426 | - | THO complex subunit 1 transcription elongatio | |
| Chr.2.734668.756644 | 2 | SNOG_08239 | 735814 | 736596 | + | Replication Fork Protection Component Swi3 | GO:0005634\|GO:0006974\|GO:0048478\|GO:0000076 |
| Chr.2.734668.756644 | 2 | SNOG_08240 | 736904 | 740926 | + | Helicase conserved C-terminal domain, DSHCT | GO:0003676\|GO:0005524\|GO:0003723\|GO:0003724\|GO:0006 |
| Chr.2.734668.756644 | 2 | SNOG_08242 | 742911 | 746393 | + | Importin-beta N-terminal domain, Cse1 | GO:0006886\|GO:0008536\|GO:0006886 |

| | | | | | | | |
|---|---|---|---|---|---|---|---|
| Chr.2.734668.756644 | 2 | SNOG_08244 | 747396 | 749334 | + | | GO:0005515 |
| Chr.2.734668.756644 | 2 | SNOG_08246 | 749954 | 756644 | + | FKBP12-rapamycin binding domain, Domain of | GO:0005515\|GO:0044877\|GO:0004674\|GO:0016301 |
| Chr.4.2141207.2169014 | 4 | SNOG_15907 | 2145758 | 2146948 | - | Eukaryotic aspartyl protease | GO:0004190\|GO:0006508 |
| Chr.4.2141207.2169014 | 4 | SNOG_15908 | 2147468 | 2148800 | + | Eukaryotic aspartyl protease | GO:0004190\|GO:0006508 |
| Chr.4.2141207.2169014 | 4 | SNOG_15915 | 2162503 | 2163068 | - | | |
| Chr.4.2141207.2169014 | 4 | SNOG_15917 | 2168175 | 2168738 | - | | GO:0005199\|GO:0031505 |
| Chr.5.1017824.1037824 | 5 | SNOG_04307 | 1034782 | 1035953 | + | | |
| Chr.5.1017824.1037824 | 5 | SNOG_04309 | 1032196 | 1032735 | + | Phenolic acid decarboxylase (PAD) | GO:0016831 |
| Chr.5.1017824.1037824 | 5 | SNOG_04312 | 1024491 | 1026128 | + | CBF/Mak21 family | GO:0042254 |
| Chr.5.1017824.1037824 | 5 | SNOG_04313 | 1023452 | 1024297 | - | Mitochondrial ATP synthase B chain precursor | GO:0000276\|GO:0015078\|GO:0015986 |
| Chr.5.1017824.1037824 | 5 | SNOG_04314 | 1021598 | 1023307 | + | | |
| Chr.5.1017824.1037824 | 5 | SNOG_04316 | 1020029 | 1021054 | + | TFIIF, beta subunit N-terminus, TFIIF, beta sub | GO:0006367\|GO:0005674\|GO:0006366\|GO:0006367 |
| Chr.5.1017824.1037824 | 5 | SNOG_04317 | 1018349 | 1019605 | - | PCI domain | GO:0008180\|GO:0005515 |
| Chr.5.1017824.1037824 | 5 | SNOG_04318 | 1017825 | 1018035 | + | Ribosomal prokaryotic L21 protein | GO:0005840 |
| Chr.6.1162062.1184015 | 6 | SNOG_07044 | 1162063 | 1162969 | + | | |
| Chr.6.1162062.1184015 | 6 | SNOG_07046 | 1166398 | 1167359 | - | EF-1 guanine nucleotide exchange domain, Eu | GO:0003746\|GO:0005853\|GO:0006414\|GO:0003746\|GO:0006 |
| Chr.6.1162062.1184015 | 6 | SNOG_07047 | 1167994 | 1170217 | + | C-terminal, D2-small domain, of ClpB protein, subfamily) | GO:0005524 |
| Chr.6.1162062.1184015 | 6 | SNOG_07051 | 1178914 | 1180437 | + | Aflatoxin regulatory protein, Fungal Zn(2)-Cys( | GO:0000981\|GO:0005634\|GO:0006355\|GO:0008270\|GO:0003 0045122 |
| Chr.8.520242.540242 | 8 | SNOG_07289 | 520243 | 520570 | - | Cyclophilin type peptidyl-prolyl cis-trans isome | GO:0000413\|GO:0003755\|GO:0003755\|GO:0003755\|GO:0006 |
| Chr.8.520242.540242 | 8 | SNOG_07290 | 520797 | 522407 | + | Putative zinc finger motif, C2HC5-type | GO:0005634\|GO:0006355\|GO:0008270 |
| Chr.8.520242.540242 | 8 | SNOG_07292 | 525556 | 525756 | + | | |
| Chr.8.520242.540242 | 8 | SNOG_07296 | 530090 | 530734 | + | | |
| Chr.8.520242.540242 | 8 | SNOG_07299 | 534972 | 537379 | + | Transient receptor potential (TRP) ion channel | |
| Chr.8.520242.540242 | 8 | SNOG_07300 | 538174 | 538525 | + | | |
| Chr.8.520242.540242 | 8 | SNOG_07303 | 539782 | 540242 | - | Bacterial low temperature requirement A prot | |
| Chr.8.520242.540242 | 8 | SNOG_30518 | 539353 | 539742 | + | | |
| Chr.9.866643.886643 | 9 | SNOG_08858 | 883727 | 885506 | + | Sugar (and other) transporter | GO:0016021\|GO:0022857\|GO:0055085 |

| | | | | | | | |
|---|---|---|---|---|---|---|---|
| Chr.9.866643.886643 | 9 | SNOG_08861 | 879254 | 880213 | - | | |
| Chr.9.866643.886643 | 9 | SNOG_08862 | 877817 | 878797 | - | | |
| Chr.9.866643.886643 | 9 | SNOG_08864 | 870495 | 872892 | - | Dipeptidyl peptidase IV (DPP IV) N-terminal reg | GO:0004252\|GO:0006508\|GO:0006508\|GO:0006508\|GO:0008 |
| Chr.9.866643.886643 | 9 | SNOG_08865 | 868738 | 869747 | - | | |
| Chr.9.866643.886643 | 9 | SNOG_08867 | 866644 | 867409 | - | Major Facilitator Superfamily | GO:0005887\|GO:0055085 |
| Chr.11.535736.555736 | 11 | SNOG_09406 | 553657 | 555375 | + | Protein kinase domain | GO:0004672\|GO:0005524\|GO:0006468\|GO:0005524\|GO:0004 |
| Chr.11.535736.555736 | 11 | SNOG_09408 | 551604 | 552709 | + | Mycolic acid cyclopropane synthetase | |
| Chr.11.535736.555736 | 11 | SNOG_09410 | 546390 | 547247 | - | | GO:0003676 |
| Chr.11.535736.555736 | 11 | SNOG_09411 | 545108 | 545831 | + | | |
| Chr.11.535736.555736 | 11 | SNOG_09412 | 541985 | 544517 | - | RhoGEF domain, Fungal N-terminal domain of | GO:0005089\|GO:0035023 |
| Chr.11.535736.555736 | 11 | SNOG_09413 | 537741 | 541709 | - | | |
| Chr.11.535736.555736 | 11 | SNOG_09414 | 535737 | 536763 | + | IQ calmodulin-binding motif, IQ calmodulin-bir | GO:0005515\|GO:0003774\|GO:0005524\|GO:0016459\|GO:0003 |
| Chr.12.200453.220453 | 12 | SNOG_03430 | 200454 | 201571 | - | Leucine rich repeat | GO:0005515 |
| Chr.12.200453.220453 | 12 | SNOG_03432 | 202901 | 203673 | - | | |
| Chr.12.200453.220453 | 12 | SNOG_03433 | 204383 | 206431 | + | | |
| Chr.12.200453.220453 | 12 | SNOG_03434 | 207713 | 208405 | + | | |
| Chr.12.200453.220453 | 12 | SNOG_03436 | 211929 | 212840 | - | | |
| Chr.12.200453.220453 | 12 | SNOG_03437 | 213639 | 214238 | - | Cysteine-rich secretory protein family | GO:0005576 |
| Chr.12.200453.220453 | 12 | SNOG_03439 | 215120 | 217571 | - | | |
| Chr.12.200453.220453 | 12 | SNOG_03440 | 219869 | 220453 | - | Sodium/calcium exchanger protein, Sodium/ca | GO:0016021\|GO:0055085 |
| Chr.15.1085013.1109854 | 15 | SNOG_14193 | 1107911 | 1109854 | - | Clr5 domain | |
| Chr.15.1085013.1109854 | 15 | SNOG_14194 | 1103845 | 1107762 | + | AAA domain | GO:0000723\|GO:0005634\|GO:0006281\|GO:0016887\|GO:0030 |
| Chr.15.1085013.1109854 | 15 | SNOG_14196 | 1101924 | 1103437 | - | PAS fold, GATA zinc finger | GO:0006355\|GO:0008270\|GO:0006355\|GO:0008270\|GO:0043 |
| Chr.15.1085013.1109854 | 15 | SNOG_14200 | 1094226 | 1095989 | - | Glycosyl hydrolase family 47 | GO:0003824\|GO:0004571\|GO:0005509\|GO:0016020 |
| Chr.15.1085013.1109854 | 15 | SNOG_14203 | 1085014 | 1089178 | + | Munc13 (mammalian uncoordinated) homolog unknown function (DUF810) | |
| Chr.18.975976.995976 | 18 | SNOG_13710 | 985746 | 990972 | - | Putative death-receptor fusion protein (DUF24 | |
| Chr.18.975976.995976 | 18 | SNOG_13711 | 983737 | 985420 | + | MBOAT, membrane-bound O-acyltransferase f | GO:0004144\|GO:0019432\|GO:0008374 |

| | | | | | | | |
|---|---|---|---|---|---|---|---|
| Chr.18.975976.995976 | 18 | SNOG_13712 | 981083 | 983236 | - | WD domain, G-beta repeat, WD domain, G-bet | GO:0005515 |
| Chr.18.975976.995976 | 18 | SNOG_13713 | 978903 | 980237 | - | Solute carrier family 35 | GO:0016021\|GO:0022857\|GO:0055085 |
| Chr.18.975976.995976 | 18 | SNOG_13714 | 977154 | 978401 | + | eIF4-gamma/eIF5/eIF2-epsilon, Domain found | GO:0003743\|GO:0006413\|GO:0005515 |
| Chr.19.954019.974019 | 19 | SNOG_11897 | 954936 | 957810 | + | Ig-like domain from next to BRCA1 gene, Zinc f | GO:0008270 |
| Chr.19.954019.974019 | 19 | SNOG_11900 | 960372 | 962829 | + | Ku70/Ku80 beta-barrel domain, Ku C terminal | GO:0003677\|GO:0006303\|GO:0000723\|GO:0003677\|GO:0003 0006310\|GO:0042162\|GO:0043564\|GO:0016817 |
| Chr.19.954019.974019 | 19 | SNOG_11902 | 963344 | 964399 | - | | |
| Chr.19.954019.974019 | 19 | SNOG_11905 | 968422 | 969491 | + | Fungal N-terminal domain of STAND proteins | |
| Chr.19.954019.974019 | 19 | SNOG_119064 | 969761 | 970862 | - | Domain found in IF2B/IF5 | GO:0003743\|GO:0006413 |
| Chr.19.954019.974019 | 19 | SNOG_11907 | 971371 | 971753 | + | | |
| Chr.19.954019.974019 | 19 | SNOG_11908 | 972659 | 973341 | + | | |
| Chr.19.954019.974019 | 19 | SNOG_11909 | 973726 | 974019 | - | | |
| Chr.20.16221.38169 | 20 | SNOG_12273 | 21468 | 23930 | - | Alpha-L-rhamnosidase N-terminal domain, Bac glycosidase domain, Bacterial alpha-L-rhamnos | GO:0003824 |
| Chr.20.16221.38169 | 20 | SNOG_12275 | 27084 | 28917 | - | | |
| Chr.20.16221.38169 | 20 | SNOG_12276 | 29700 | 30188 | + | | |
| Chr.20.16221.38169 | 20 | SNOG_12277 | 30422 | 31284 | - | | GO:0048037 |
| Chr.20.16221.38169 | 20 | SNOG_12278 | 31770 | 34091 | - | Fungal specific transcription factor domain | GO:0000981\|GO:0005634\|GO:0006355\|GO:0008270\|GO:0003 0008270 |
| Chr.20.16221.38169 | 20 | SNOG_12279 | 34750 | 37596 | + | | GO:0046872 |
| Chr.20.16221.38169 | 20 | SNOG_30828 | 18670 | 18977 | + | | |
| Chr.20.16221.38169 | 20 | SNOG_30829 | 25683 | 26414 | - | | |
| Chr.23.409903.429903 | 23 | SNOG_16288 | 421904 | 423559 | + | | |
| Chr.23.409903.429903 | 23 | SNOG_16291 | 416929 | 417753 | - | | |
| Chr.23.409903.429903 | 23 | SNOG_16292 | 415162 | 416172 | + | | |

**Supplementary Table 4.** Gene ontology (GO) term enrichment summary for the genetic groups of *Parastagonospora nodorum*.

| Genetic group | Test direction | Ontology category | GO Term | Enrichment p-value | Odds Ratio | Expected GO term count | Effective GO term count | Total genes per GO | Term |
|---|---|---|---|---|---|---|---|---|---|
| 1 | Over represented | Biological process | GO:0000723 | 0.0023 | 35.774 | 0.074 | 2 | 10 | telomere maintenance |
| | | | GO:0006413 | 0.0111 | 14.271 | 0.162 | 2 | 22 | translational initiation |
| | | | GO:0065007 | 0.0152 | 2.589 | 4.784 | 10 | 649 | biological regulation |
| | | | GO:0009889 | 0.0153 | 3.064 | 2.698 | 7 | 366 | regulation of biosynthetic process |
| | | | GO:0034645 | 0.0200 | 2.386 | 5.757 | 11 | 781 | cellular macromolecule biosynthetic process |
| | | | GO:0044260 | 0.0226 | 2.983 | 3.000 | 7 | 504 | cellular macromolecule metabolic process |
| | | | GO:0044238 | 0.0316 | 2.113 | 17.241 | 23 | 2339 | primary metabolic process |
| | | | GO:0034641 | 0.0327 | 2.049 | 9.582 | 15 | 1300 | cellular nitrogen compound metabolic process |
| | | | GO:1901576 | 0.0354 | 2.070 | 7.924 | 13 | 1075 | organic substance biosynthetic process |
| | | | GO:0042128 | 0.0363 | 34.688 | 0.037 | 1 | 5 | nitrate assimilation |
| | | | GO:0045229 | 0.0363 | 34.688 | 0.037 | 1 | 5 | external encapsulating structure organization |
| | | | GO:2001057 | 0.0363 | 34.688 | 0.037 | 1 | 5 | reactive nitrogen species metabolic process |
| | | | GO:0019219 | 0.0380 | 2.690 | 2.553 | 6 | 357 | regulation of nucleobase-containing compound metabolic process |
| | | | GO:0006355 | 0.0419 | 2.616 | 2.609 | 6 | 354 | regulation of transcription, DNA-templated |
| | | | GO:2001141 | 0.0419 | 2.616 | 2.609 | 6 | 354 | regulation of RNA biosynthetic process |
| | | | GO:0034654 | 0.0470 | 2.247 | 4.143 | 8 | 562 | nucleobase-containing compound biosynthetic process |
| | | Celular component | GO:0016459 | 0.0493 | 25.726 | 0.050 | 1 | 5 | myosin complex |
| | | Molecular function | GO:0017016 | 0.0127 | 13.107 | 0.173 | 2 | 23 | Ras GTPase binding |
| | | | GO:0004190 | 0.0149 | 11.963 | 0.188 | 2 | 25 | aspartic-type endopeptidase activity |
| | | | GO:0003743 | 0.0211 | 9.818 | 0.226 | 2 | 30 | translation initiation factor activity |
| | | | GO:0051020 | 0.0282 | 8.323 | 0.263 | 2 | 35 | GTPase binding |
| | | | GO:0051015 | 0.0371 | 33.703 | 0.038 | 1 | 5 | actin filament binding |

| | | | | | | | | | |
|---|---|---|---|---|---|---|---|---|---|
| | | | GO:0003951 | 0.0371 | 33.703 | 0.038 | 1 | 5 | NAD+ kinase activity |
| | | | GO:0008270 | 0.0428 | 2.363 | 3.290 | 7 | 437 | zinc ion binding |
| | | | GO:0008374 | 0.0443 | 26.958 | 0.045 | 1 | 6 | O-acyltransferase activity |
| | | | GO:0003746 | 0.0443 | 26.958 | 0.045 | 1 | 6 | translation elongation factor activity |
| | | | GO:0140096 | 0.0444 | 2.215 | 4.042 | 8 | 537 | catalytic activity, acting on a protein |
| | Under represented | Biological process | GO:0055114 | 0.0118 | 0.142 | 5.926 | 1 | 804 | oxidation-reduction process |
| | | Molecular function | GO:0016491 | 0.0039 | 0.120 | 7.121 | 1 | 946 | oxidoreductase activity |
| | | | GO:0048037 | 0.0379 | 0.187 | 4.833 | 1 | 642 | cofactor binding |
| | | | GO:0003824 | 0.0458 | 0.573 | 25.045 | 19 | 3327 | catalytic activity |
| 2 | Over represented | Molecular function | GO:0005515 | 0.008540399 | 2.310 | 7.949 | 15 | 968 | protein binding |
| 3 | Over represented | Molecular function | GO:0000981 | 0.018222804 | 3.110 | 2.114 | 6 | 213 | DNA-binding transcription factor activity, RNA polymerase II-specific |
| | | | GO:0048037 | 0.04826314 | 1.912 | 6.371 | 11 | 642 | cofactor binding |
